# Prostaglandin in the ventromedial hypothalamus regulates peripheral glucose metabolism

**DOI:** 10.1101/2020.05.18.056374

**Authors:** Ming-Liang Lee, Hirokazu Matsunaga, Yuki Sugiura, Takahiro Hayasaka, Izumi Yamamoto, Daigo Imoto, Makoto Suematsu, Norifumi Iijima, Kazuhiro Kimura, Sabrina Diano, Chitoku Toda

## Abstract

The hypothalamus plays a central role in monitoring and regulating systemic glucose metabolism. The brain is enriched with phospholipids containing poly-unsaturated fatty acids, which are biologically active in physiological regulation. Here, we show that intraperitoneal glucose injection induced changes in hypothalamic distribution and amount of phospholipids, especially arachidonic-acid-containing phospholipids, that were then metabolized to produce prostaglandins. Knockdown of cytosolic phospholipase A2 (cPLA2), a key enzyme for generating arachidonic acid from phospholipids, in the hypothalamic ventromedial nucleus (VMH), lowered insulin sensitivity in muscles during regular chow diet (RCD) feeding. Conversely, the down-regulation of glucose metabolism by high fat diet (HFD) feeding was improved by knockdown of cPLA2 in the VMH through changing hepatic insulin sensitivity and hypothalamic inflammation. Our data suggest that cPLA2-mediated hypothalamic phospholipid metabolism is critical for controlling systemic glucose metabolism during RCD, while continuous activation of the same pathway to produce prostaglandins during HFD deteriorates glucose metabolism.

## Introduction

Recent evidence from animal models indicates that the brain plays a critical role in the systemic regulation of glucose metabolism^1,2^. Neurons in the hypothalamus integrate hormonal and nutritional information and maintain glucose homeostasis by controlling metabolism in peripheral tissues. Numerous brain regions have been reported to maintain whole-body glucose homeostasis^3–5^. In particular, the ventromedial nucleus of the hypothalamus (VMH) and arcuate nucleus of the hypothalamus (ARC) are critical nuclei for the glucose metabolism. Obesity attenuates the function of these nuclei and promotes type II diabetes via hypothalamic inflammation^6^. However, the hypothalamic mechanism that regulates systemic glucose metabolism is not fully understood.

The VMH has important roles in regulating glucose metabolism in peripheral tissues^7^, and the majority of neurons in the VMH express steroidogenic factor 1 (Sf1). Photoactivation of Sf1 neurons increases hepatic glucose production (HGP)^4,8^ and simultaneously enhances insulin sensitivities in the liver, muscle and brown adipose tissue (BAT)^9^. Leptin regulates the neuronal activity of Sf1 neurons, and increases glucose utilization and insulin sensitivity in peripheral tissues^10–12^. There are two main neuronal populations in the ARC, the orexigenic agouti-related peptide (AgRP) neurons, which co-express neuropeptide Y (NPY), and the anorexigenic proopiomelanocortin (POMC) neurons^13^. POMC and AgRP neurons control HGP in opposite ways, i.e., activation of POMC increases insulin sensitivity in the liver, while AgRP activation decreases liver insulin sensitivity^14,15^. Hormones, including insulin, leptin and ghrelin, regulate glucose metabolism by changing the activities of these neurons and their gene expression. In addition, subpopulations of Sf1, POMC and AgRP neurons are also activated (glucose excited neurons) or inhibited (glucose inhibited neurons) by glucose^16^.

Fatty acids regulate the activities of hypothalamic neurons^16^ and the lipid metabolism within the hypothalamus plays important roles in energy balance and glucose metabolism^17,18^. Phospholipids with biologically active polyunsaturated fatty acids (PUFAs), including phosphatidyl-inositol (PI), phosphatidyl-ethanolamine (PE) and phosphatidyl-serine (PS), are abundantly found in the brain^19^. Some membrane phospholipids generate free PUFAs to regulate physiological functions of the brain. For example, phospholipase A2 (PLA2) preferentially generates arachidonic acid (AA) from phospholipids^20^. AA plays roles in several physiological functions, including thermogenesis in BAT and increasing blood glucose levels^21^. AA is also the precursor for eicosanoids such as, prostaglandin and hydroxyeicosatetraenoic acid (HETE). Other PUFA, such as oleic acid (OA), modulates activities of nutrient-sensing neurons to regulate insulin secretion^22^, and intracerebroventricular injection of OA enhances insulin sensitivity in the liver^23^. However, the distributions of FAs, PUFAs and PUFA-containing phospholipids in the hypothalamus and their roles in whole-body glucose metabolism are not clearly understood.

Here, we explore hypothalamic lipid metabolism in the regulation of systemic glucose homeostasis and its potential role in the development of diabetes using imaging mass spectrometry (IMS). We found that glucose injection in mice fed on a regular chow diet induces a decrease in phospholipids containing AA, which is mediated by cytosolic phospholipase A2 (cPLA2). Prostaglandins produced by phospholipids in the hypothalamus activates VMH neurons and increases insulin sensitivity in skeletal muscles. However, hypothalamic cPLA2-mediated prostaglandin production is enhanced by high-fat-diet and induces neuroinflammation, and blockage of this enzyme confers resistance to developing diabetes.

## Results

### Hyperglycemia decreases phospholipids and produces prostaglandins in the hypothalamus

To determine the distribution of FAs and phospholipids, hypothalamic slices were examined by IMS (Fig. 1). The signal intensities of palmitic acid (PA), stearic acid (SA), AA and docosahexaenoic acid (DHA) were high around the third ventricle and the ventro-lateral region of the hypothalamus in C57BL/6J mice (Fig. 1a). Similar distributions of phospholipids, such as phosphatidyl-ethanolamine (PE) (18:0/20:4), phosphatidyl-inositol (PI) (18:0/20:4) and PI (18:1/20:4) were also observed. However, signal intensities for phosphatidyl-serine (PS) (18:0/20:4) were low around the third ventricle while PS (18:0/22:6) distribution was ubiquitously observed (Fig. 1b). We then measured hypothalamic lipids in mice that received an intraperitoneal (i.p.) glucose injection. Signal intensities for PI (18:0/20:4), PI (18:1/20:4) and PE (18:0/20:4) were significantly decreased in the VMH after glucose administration (Fig. 1c,d). Similarly, the signal intensities for PI (18:0/20:4), PI (18:1/20:4), PE (18:0/20:4) and PS (18:0/16:0) were decreased in the ARC after glucose injection (Fig. 1c,e). Hydrolysis of these phospholipids generates FAs, including AA (20:4), oleic acid (OA) (18:1), PA (16:0) or SA (18:0). However, the signal intensities of the four fatty acids were not changed after glucose injection, neither in the VMH nor ARC (Fig. 1f-h).

**Figure 1.**
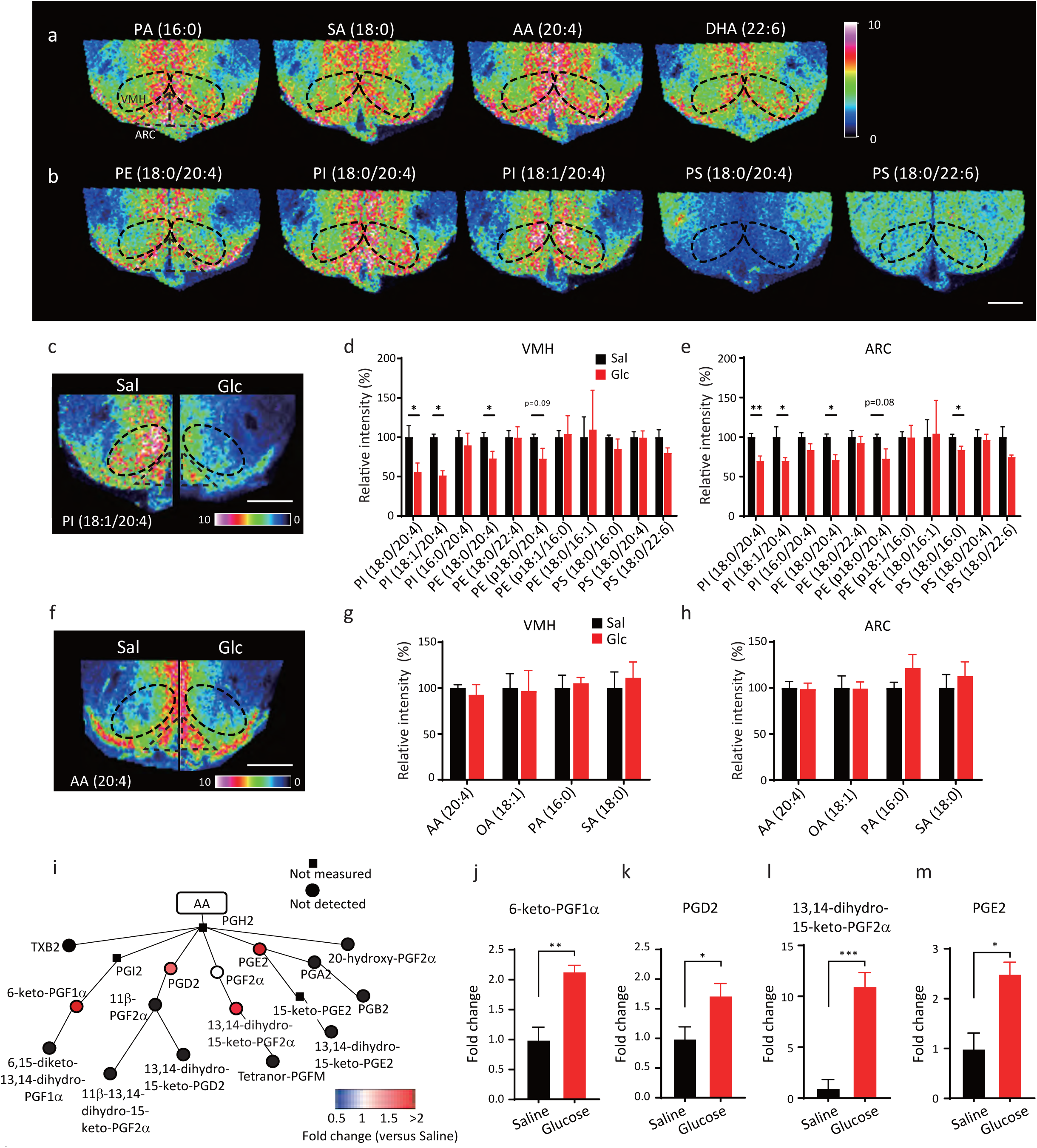
Hyperglycemia increases prostaglandin production derived from phospholipids. **a**,**b**, Representative results of imaging mass spectrometry (IMS) showing distributions of hypothalamic fatty acids (**a**) and phospholipids (**b**) from untreated RCD-fed mice. The dashed black line shows the position of the VMH. Scale bar: 500 μm. **c-h**, Distributions of phospholipids and fatty acids in the hypothalamus 30 min after injection of saline or glucose (2 g/kg). **c**,**f**, Representative results of IMS on hypothalamic phosphatidylinositol (PI) (18:1/20:4) (**c**) and arachidonic acid (AA) (**f**) from mice i.p. injected with saline (left half) or glucose (right half). Scale bar: 500 μm. **d**,**e**, Relative intensities of phospholipids in the VMH (**d**) and ARC (**e**) after injection with saline (n=5) or glucose (n=4). **g**,**h**, Relative intensities of fatty acids in the VMH (**g**) and ARC (**h**) after injection with saline (n=5) or glucose (n=4). **i-m**, LC-MS results showing the effects of glucose injection on AA metabolites in the whole hypothalamus. (**i**) Relative amounts of hypothalamic prostaglandins mediated by cyclooxygenase. (**j**) 6-keto-PGF1α, (**k**) PGD2, (**l**) 13,14-dihydro-15-keto-PGF2α and (**m**) PGE2 were increased by glucose injection. n=5 in each experimental group. **d-h** and **j-l** represent the mean ± SEM; * = p<0.05; ** = p<0.01; *** = p<0.001. **i**, represents the mean fold change in color.

AA is the source of eicosanoids and AA metabolism is catalyzed by enzymes such as cyclooxygenase (COX), lipoxygenase and cytochrome P450. To elucidate if glucose injection increased AA metabolism, we examined the effect of glucose injection on eicosanoid production in the whole hypothalamus using a liquid chromatography–mass spectrometry (LC-MS). Compared with saline, glucose injection increased COX-mediated hypothalamic production of prostaglandins, including 6-keto-PGF1α, PGD2, 13,14-dihydro-15-keto-PGF2α and PGE2 (Fig. 1i-m). Lipoxygenase-mediated production of 12-HETE was increased in glucose-injected mice (Fig. S1a). However, most of the lipoxygenase- and cytochrome P450-mediated production of HETEs and EETs was not detected or changed by glucose injection (Fig. S1a,b). Thus, the data suggest that increased glucose levels decreases AA-containing phospholipids to produce prostaglandins.

### Blocking the PLA2-mediated pathway in the hypothalamus impairs systemic glucose metabolism

PLA2 is the primary enzyme that generates AA from phospholipids^24^. We then investigated the role of PLA2-mediated phospholipid utilization in glucose metabolism during acute hyperglycemia after an intrahypothalamic administration of methyl arachidonyl fluorophosphonate (MAFP), a PLA2 inhibitor. MAFP-injected mice showed decreased glucose tolerance compared to vehicle-injected mice and no changes in circulating insulin levels were observed (Fig. 2a,b). Hypothalamic injection of indomethacin, an inhibitor of COX1/2, also impaired glucose tolerance, suggesting that prostaglandins regulate hypothalamic function to decrease blood glucose levels (Fig. 2c,d). However, intra-hypothalamic injection of phospholipase C (PLC) inhibitor, U73122, or IP3 receptor antagonist, xestospondin 2, did not affect glucose tolerance (Fig. S2). Thus, our results suggest that the PLA2-mediated AA release and production of prostaglandins by COX1/2 in the hypothalamus, but not PLC-IP3 pathway, play a role in regulating glucose metabolism during acute hyperglycemia.

**Figure 2.**
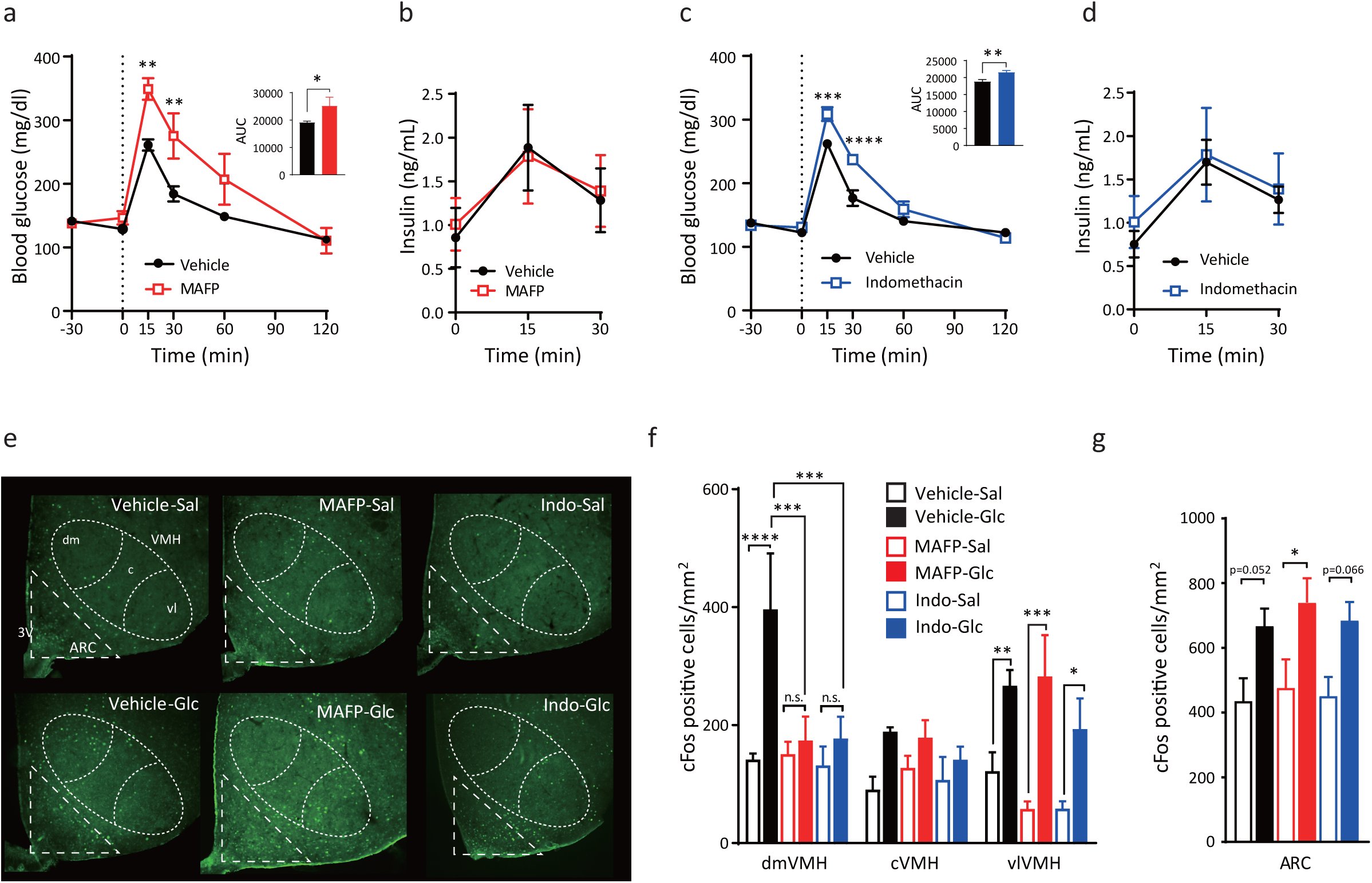
Hypothalamic PLA2- and COX-mediated AA metabolism regulates systemic glucose tolerance and modulates glucose responsiveness in the dmVMH. **a**, Glucose tolerance test (GTT) (0–120 min) after intra-hypothalamic injection (−30 min) of MAFP (n=7) or vehicle (n=7). **b**, Blood insulin concentration of MAFP (n=7) or vehicle (n=7) injected mice during GTT. **c**, GTT (0–120 min) after intra-hypothalamic injection (−30 min) of indomethacin (n=7) or vehicle (n=7). **d**, Blood insulin concentration of indomethacin (n=7) or saline (n=7) injected mice during GTT. **e**, Representative micrographs showing immunofluorescent cFos staining in the hypothalamus of saline (upper panels) or glucose (lower panels) injected mice after i.c.v. injection of PBS, MAFP or indomethacin (indo). Scale bar: 500 μm. dm: dorsomedial, c: central, vl: ventrolateral part of the VMH. **f**,**g**, Quantification of cFos expression in the dorsomedial (dmVMH), central (cVMH) and ventrolateral (vlVMH) subregions of the VMH (**f**) and ARC (**g**) from mice injected with saline or glucose after i.c.v. injection of PBS, MAFP or indomethacin (n=3 in each experimental group). All data represent the mean ± SEM; * = p<0.05; ** = p<0.01; *** = p<0.001; **** = p<0.0001.

### PLA2-mediated production of prostaglandin is necessary for the responsiveness of the VMH to glucose

To understand the role of PLA2 in controlling glucose metabolism, we examined the effect of PLA2 inhibitors on hypothalamic neuronal activation by cFos expression. Vehicle, MAFP or indomethacin were injected intracerebroventricularly (i.c.v.) 30 minutes prior to i.p. injection of either saline or glucose in fasted mice. In i.c.v. vehicle-injected mice, glucose administration increased cFos-positive cells in the dorsomedial (dm) and ventrolateral (vl) regions of the VMH and in the ARC (Fig. 2e-g). In i.c.v. MAFP-injected mice, glucose did not alter the number of cFos-positive neurons in the dmVMH (Fig. 2f). An increase in cFos-positive neurons after glucose injection was still detected in the vlVMH and ARC compared with saline injected mice (Fig. 2f,g). Similar results were observed in i.c.v. indomethacin-injected mice after an i.p. injection of glucose (Fig. 2f,g). Taken together, these data showing that both MAFP and indomethacin block neuronal activation during acute hyperglycemia in the dmVMH, indicates that metabolites of phospholipid-derived prostaglandins regulate glucose responsiveness of neurons in the dmVMH, while glucose activates neurons in the vlVMH and ARC independently of PLA2 and COX1/2.

### Knockdown of *pla2g4a* in Sf1 neurons impairs glucose metabolism in regular chow diet feeding

Next, to explore the role of PLA2 in VMH neurons, short hairpin RNA (shRNA) against *pla2g4a*, a gene encoding cytosolic PLA2 (cPLA2), which has specificity for *sn*-2 arachidonic acid and a role in eicosanoid production^20^, was transfected to the VMH through an adeno-associated virus (AAV) cre-recombinase (cre)-dependent in Sf1-cre mice (Fig. S3a,b). Expression of *pla2g4a* mRNA was significantly decreased in the VMH of Sf1-cre mice injected with AAV-DIO-shRNA (cPLA2KD^Sf1^) compared with AAV-DIO-GFP (GFP^Sf1^)-injected mice (Fig. S3c). Although knockdown of *pla2g4a* did not influence body weight or the weight of adipose tissues, muscle and liver (Fig. S3d), cPLA2KD^Sf1^ mice displayed decreased glucose tolerance and insulin sensitivity compared with GFP^Sf1^ mice (Fig. 3a,b). To rule out the involvement of astrocytic cPLA2, AAV-GFAP-Cre and AAV-DIO-shRNA against *pla2g4a* were co-injected into the hypothalamus to knock down the expression of *pla2g4a* in hypothalamic astrocytes (Fig. S4a,b). The knockdown of cPLA2 in astrocytes did not alter glucose metabolism, insulin sensitivity or body weight compared with control mice (Fig. S4c-e). Thus, our data suggest that cPLA2 in Sf1 neurons, not astrocytes, regulates peripheral glucose metabolism.

**Figure 3.**
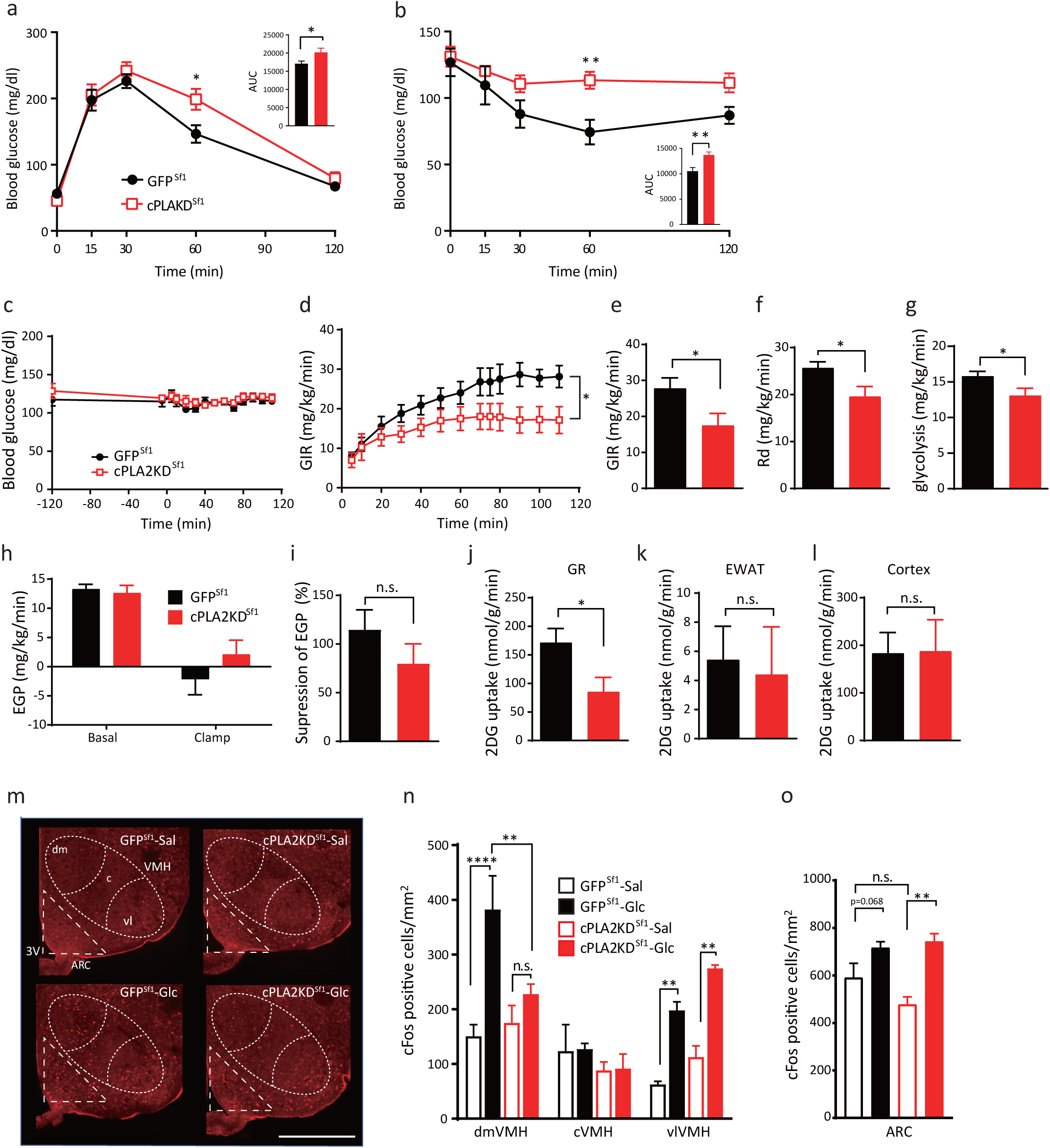
Knockdown of Sf1-neuronal *pla2g4a* impairs systemic glucose metabolism. **a**, Glucose tolerance test in cPLA2KD ^Sf1^ (n=6) and GFP^Sf1^ mice (n=6). **b**, Insulin tolerance test in cPLA2KD^Sf1^ (n=6) and GFP^Sf1^ mice (n=6). **c-l**, Hyperinsulinemic–euglycemic clamp studies in cPLA2KD^Sf1^ and GFP^Sf1^ mice. **c**, Blood glucose levels during hyperinsulinemic–euglycemic clamp studies in cPLA2KD^Sf1^ or GFP^Sf1^ mice. **d**, The glucose infusion rate (GIR) required to maintain euglycemia during the clamp period in cPLA2KD^Sf1^ (n=7) or GFP^Sf1^ mice (n=7). **e**, The average GIR between 75 and 115 min in cPLA2KD^Sf1^ (n=7) or GFP^Sf1^ mice (n=7). **f**, The rate of glucose disappearance (Rd) during the clamp period, which represents whole-body glucose utilization. **g**, The rates of whole-body glycolysis in cPLA2KD^Sf1^ (n=7) or GFP^Sf1^ mice (n=7). **h**, Endogenous glucose production (EGP) during both the basal and clamp periods in cPLA2KD^Sf1^ (n=7) or GFP^Sf1^ (n=7). **i**, Insulin-induced suppression of EGP in cPLA2KD^Sf1^ (n=7) or GFP^Sf1^ (n=7). **j-l**, Graphs showing 2-[^14^C]-Deoxy-D-Glucose uptake in red portions of the gastrocnemius (GR; **j**), white adipocyte (EWAT; **k**) and brain (cortex; **l**) during the clamp period in cPLA2KD^Sf1^ (n=7) or GFP^Sf1^ mice (n=7). **m**, Representative micrographs showing immunofluorescent cFos staining in the hypothalamus of cPLA2KD^Sf1^ and GFP^Sf1^ mice after saline or glucose injection (3 g/kg). Scale bar: 500 μm. **n**,**o**, Quantification of cFos expression in the dmVMH, cVMH, vlVMH and ARC of cPLA2KD^Sf1^ or GFP^Sf1^ mice after saline (n=3) or glucose (n=3) injection (3 g/kg). All data represent the mean ± SEM; * = p<0.05; ** = p<0.01; *** = p<0.001; **** = p<0.0001.

To further investigate the role of cPLA2 in Sf1 neurons in glucose metabolism, we next performed hyperinsulinemic–euglycemic clamp studies. cPLA2KD^Sf1^ mice showed a lower glucose infusion rate (GIR) to maintain euglycemia compared with GFP^Sf1^ mice (Fig. 3c-e). The rate of disappearance (Rd) and glycolysis were also lower in cPLA2KD^Sf1^ mice compared with GFP^Sf1^ mice (Fig. 3f,g). However, endogenous glucose production (EGP) was not different between the two experimental groups (Fig. 3h,i), suggesting that glucose utilization, rather than EGP, was impaired in cPLA2KD^Sf1^ mice. In agreement with this, cPLA2KD^Sf1^ mice showed decreased 2DG uptake in the red part of gastrocnemius muscle (GR) compared with control mice (Fig. 3j). 2DG uptake in white adipose tissue (WAT) and the brain (cortex) were similar between groups (Fig. 3k,l).

To assess changes in neuronal activation, we next analyzed cFos expression in cPLA2KD^Sf1^ mice compared with controls. Glucose-induced cFos expression in the dmVMH of cPLA2KD^Sf1^ mice was blunted compared with glucose-injected control mice (Fig. 3m,n). The glucose-induced cFos expression in either vlVMH or ARC was not changed after the knockdown of cPLA2 (Fig. 3m,o).

Taken together, our data suggest that cPLA2-mediated prostaglandin production regulates glucose-induced activation of dmVMH neurons to control insulin sensitivity in muscle.

### High fat diet decreases AA-containing phospholipids and produces prostaglandins in the hypothalamus

High-fat-diet (HFD) induces inflammation and impairs hypothalamic functions^25^. Long chain fatty acyl CoA, a proinflammatory signal, accumulates in the hypothalamus during HFD feeding^26^. Thus, we examined the effect of HFD on lipid distribution in the hypothalamus. In mice fed an HFD for 8 weeks, the signal intensities for FAs, including AA, were greater in the ARC but not the VMH than those observed in control mice fed a RCD (Fig. 4a-c). However, signal intensities for phospholipids in the hypothalamus were lower in HFD-fed mice (Fig. 4d-f). In both the VMH and ARC, the signal intensities for PI (18:0/20:4), PI (18:1/20:4), PE (18:0/20:4), PE (p18:0/20:4) and PS (18:0/22:6) were significantly decreased in HFD-fed mice (Fig. 4e,f). Because PLA2 generates AA from these phospholipids to regulate cellular activities, we next analyzed the activity of hypothalamic cPLA2 and found that cPLA2 activity was higher in HFD-fed mice compared with RCD-fed mice (Fig. 4g). However, the activity of secretory phospholipase A2 (sPLA2) remained similar between RCD- and HFD-fed mice (Fig. 4h).

**Figure 4.**
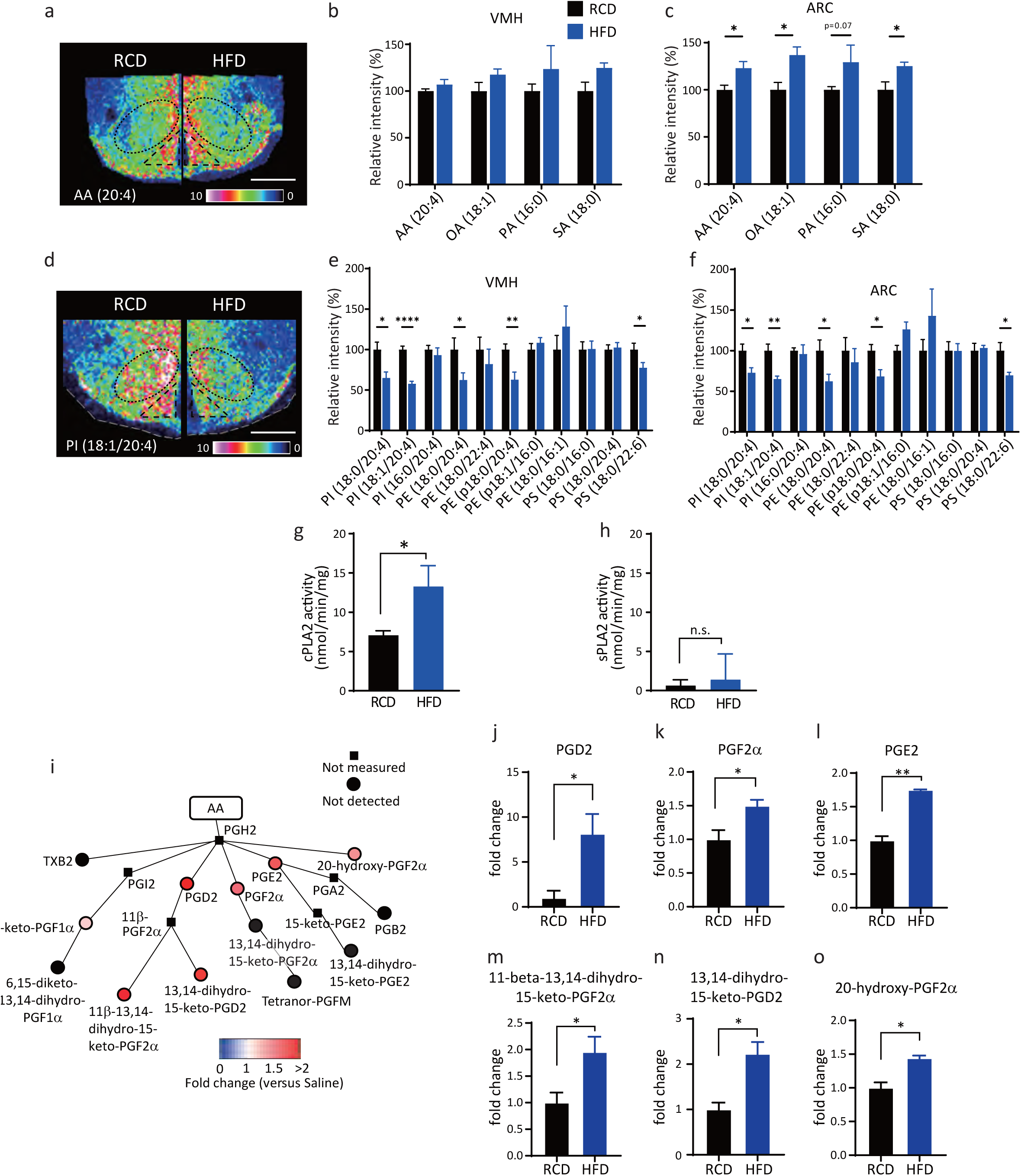
HFD feeding increases prostaglandin production derived from phospholipids. **a-f**, Distributions of fatty acids and phospholipids in the hypothalamus in RCD- or HFD-fed mice for 8 weeks. **a**,**d**, Representative results of IMS on hypothalamic arachidonic acid (AA) (**a**) and PI (18:1/20:4) (**d**) from RCD-fed mice (left) or HFD-fed mice (right). Scale bar: 500 μm. **b**,**c**, Relative intensities of fatty acids in the VMH (**b**) or ARC (**c**) of RCD- (n=5) or HFD-fed mice (n=4). **e**,**f**, Relative intensities of phospholipids in the VMH (**e**) or ARC (**f**) of RCD- (n=5) or HFD-fed (n=4) mice. **g**,**h**, Enzymatic activity of hypothalamic cPLA2 (**g**) and sPLA (**h**) in RCD- (n=5) or HFD-fed (n=5) mice. **i**, Relative amounts of prostaglandins in the hypothalamus after 8 weeks in HFD-fed mice (n=3) compared with those of RCD-fed mice (n=3). **j-o**, Bar graphs showing COX-mediated production of (**j**) PGD2, (**k**) PGF2α, (**l**) PGE2, (**m**) 11-beta-13,14-dihydro-15-keto-PGF2α, (**n**) 13,14-dihydro-15-keto-PGD2 and (**o**) 20-hydroxy-PGF2α in 8 weeks of HFD-fed mice (n=3) compared with RCD-fed mice (n=3). All data represent the mean ± SEM; * = p<0.05; ** = p<0.01; *** = p<0.001; **** = p<0.0001.

We next explored the effect of HFD on the production of eicosanoids in the hypothalamus by LC-MS (Fig. 4i and Fig. S5). In HFD-fed mice, COX-mediated production of prostaglandins, including PGD2, PGF2α, PGE2, 11-beta-13,14-dihydro-15-keto-PGF2α, 13,14-dihydro-15-keto-PGD2 and 20-hydroxy-PGF2α, was increased compared with RCD-fed mice (Fig. 4j-o). Only 12-HETE, an eicosanoid mediated by lipoxygenase, was significantly increased after HFD feeding (Fig. S5).

### Knockdown of *pla2g4a* improves HFD-induced impairment of glucose metabolism and recovers glucose responsiveness of the vlVMH and ARC to hyperglycemia

To understand the role of HFD-induced activation of hypothalamic cPLA2, we fed cPLA2KD^Sf1^ mice with HFD and examined the role of cPLA2 on glucose metabolism. Body weight and tissue weight of HFD-fed cPLA2KD^Sf1^ mice (cPLA2KD^Sf1^-HFD) were comparable to those of HFD-fed control GFP^Sf1^ mice (GFP^Sf1^-HFD) (Fig. 5a and Fig. S6). Unlike RCD-fed mice, knockdown of cPLA2 in Sf1 neurons increased glucose tolerance (Fig. 5b). However, insulin tolerance test showed no difference between groups (Fig. 5c). In GFP^Sf1^-HFD mice, no significant changes in cFos-positive neurons in the VMH or ARC were observed after glucose injection compared with saline injection (Fig. 5d-f). However, in cPLA2KD^Sf1^-HFD, the number of cFos-positive neurons were significantly higher in the vlVMH and ARC after glucose injection compared with saline injection, suggesting that cPLA2 knockdown improved neuronal responsiveness to glucose (Fig. 5d-f).

**Figure 5.**
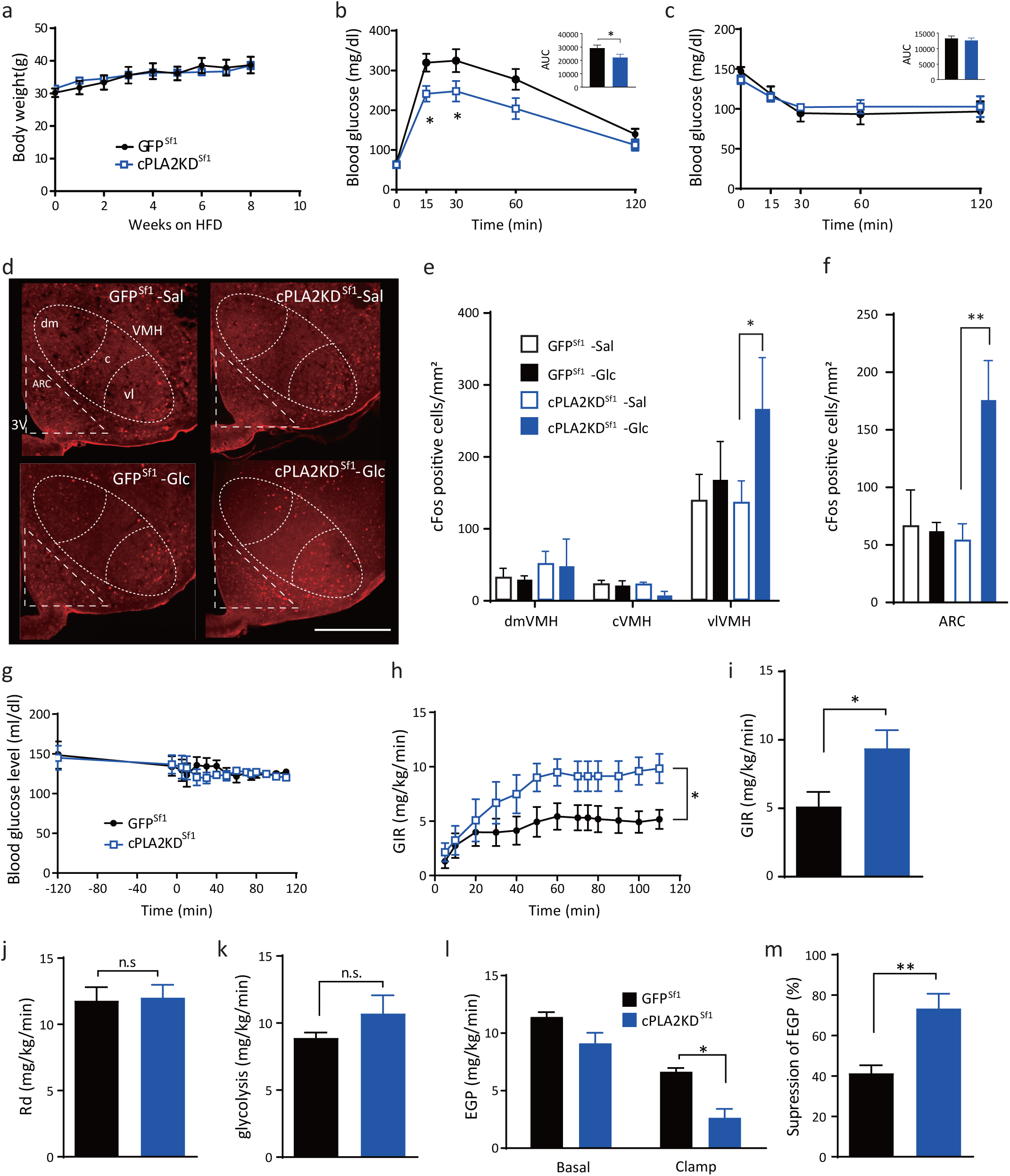
Knockdown of cPLA2 improves HFD-induced impairment of glucose metabolism. **a**, Body weight change in cPLA2KD^Sf1^ mice (n=12) and GFP^Sf1^ mice (n=10). **b**, Glucose tolerance test on cPLA2KD^Sf1^ mice (n=12) and GFP^Sf1^ mice (n=10). **c**, Insulin tolerance test on cPLA2KD^Sf1^ (n=8) mice and GFP^Sf1^ mice (n=6) during 8 weeks of HFD feeding. **d**, Representative micrographs showing immunofluorescent cFos staining in the hypothalamus of HFD-fed cPLA2KD^Sf1^ and GFP^Sf1^ mice after saline or glucose injection. Scale bar: 500 μm. **e**,**f**, Quantification of cFos expression in the dmVMH, cVMH, vlVMH (**e**), and ARC (**f**) of HFD-fed cPLA2KD^Sf1^ or GFP^Sf1^ mice after saline or glucose injection (n=3–5 in each experimental group). **g-m**, Hyperinsulinemic–euglycemic clamp studies in HFD-fed cPLA2KD^Sf1^ (n=7) or GFP^Sf1^ (n=7) mice. **g**, Blood glucose levels during hyperinsulinemic–euglycemic clamp studies in HFD-fed cPLA2KD^Sf1^ (n=7) or GFP^Sf1^ (n=7) mice. **h**, The glucose infusion rate (GIR) required to maintain euglycemia during the clamp period in cPLA2KD^Sf1^ (n=7) or GFP^Sf1^ (n=7) mice. **i**, The average GIR between 75 and 115 min in cPLA2KD^Sf1^ or GFP^Sf1^ mice. **j**, The rate of glucose disappearance (Rd) during the clamp period, which represents whole body glucose utilization. **k**, The rates of whole-body glycolysis in cPLA2KD^Sf1^ or GFP^Sf1^ mice. **l**, Endogenous glucose production (EGP) during both basal and clamp periods in cPLA2KD^Sf1^ or GFP^Sf1^ mice. **m**, The percent-suppression levels of EGP induced by insulin infusion in cPLA2KD^Sf1^ (n=7) or GFP^Sf1^ (n=7) mice. All data represent the mean ± SEM; * = p<0.05; ** = p<0.01.

To understand the role of hypothalamic cPLA2 on glucose metabolism in HFD-fed mice, we performed hyperinsulinemic–euglycemic clamping (Fig. 5g-m). To maintain euglycemia, GIR was significantly higher in cPLA2KD^Sf1^-HFD than in GFP^Sf1^-HFD (Fig. 5g-i). Unlike RCD-fed mice, glucose utilization (Rd and glycolysis) in HFD-fed mice was not altered by knocking down cPLA2 (Fig. 5j,k). In contrast, insulin inhibition of EGP was stronger during the clamp period in cPLA2KD^Sf1^-HFD mice than in GFP^Sf1^-HFD (Fig. 5l,m). These results suggest that cPLA2 in the VMH has a deteriorative role in the glucose responsiveness of the vlVMH/ARC and attenuates hepatic insulin sensitivity during HFD-induced obesity. These data suggest that the role of cPLA2 and the mechanism to change glucose metabolism and neuronal activity by prostaglandins are different between HFD and RCD.

### Knockdown of cPLA2 in Sf1 neurons prevents hypothalamic inflammation

We examined the effect of cPLA2 knockdown on hypothalamic inflammation, which attenuates neuronal functions, in HFD-fed mice. The mice were fed with a RCD or HFD for 8 weeks, and inflammation was measured by comparing the number of microglia and the astrocyte population. For this experiment, we used Iba1 and GFAP as markers of microglia and astrocytes, respectively (Fig. 6). The number of Iba1 cells was increased in the ARC but not in the VMH after mice were fed with an HFD (Fig. 6a-c). The numbers of Iba1 cells in the ARC, but not the VMH, were significantly decreased in a cPLA2KD^Sf1^-HFD compared with a GFP^Sf1^-HFD (Fig. 6b,c). HFD also increased the number of GFAP cells in the ARC, but not in the VMH, compared with mice fed a RCD (Fig. 6d-f). In the ARC, the number of GFAP-positive cells decreased in the cPLA2KD^Sf1^-HFD mice compared with GFP^Sf1^-HFD mice (Fig. 6e,f), suggesting that cPLA2 in the VMH has an influence on inflammation in the ARC.

**Figure 6.**
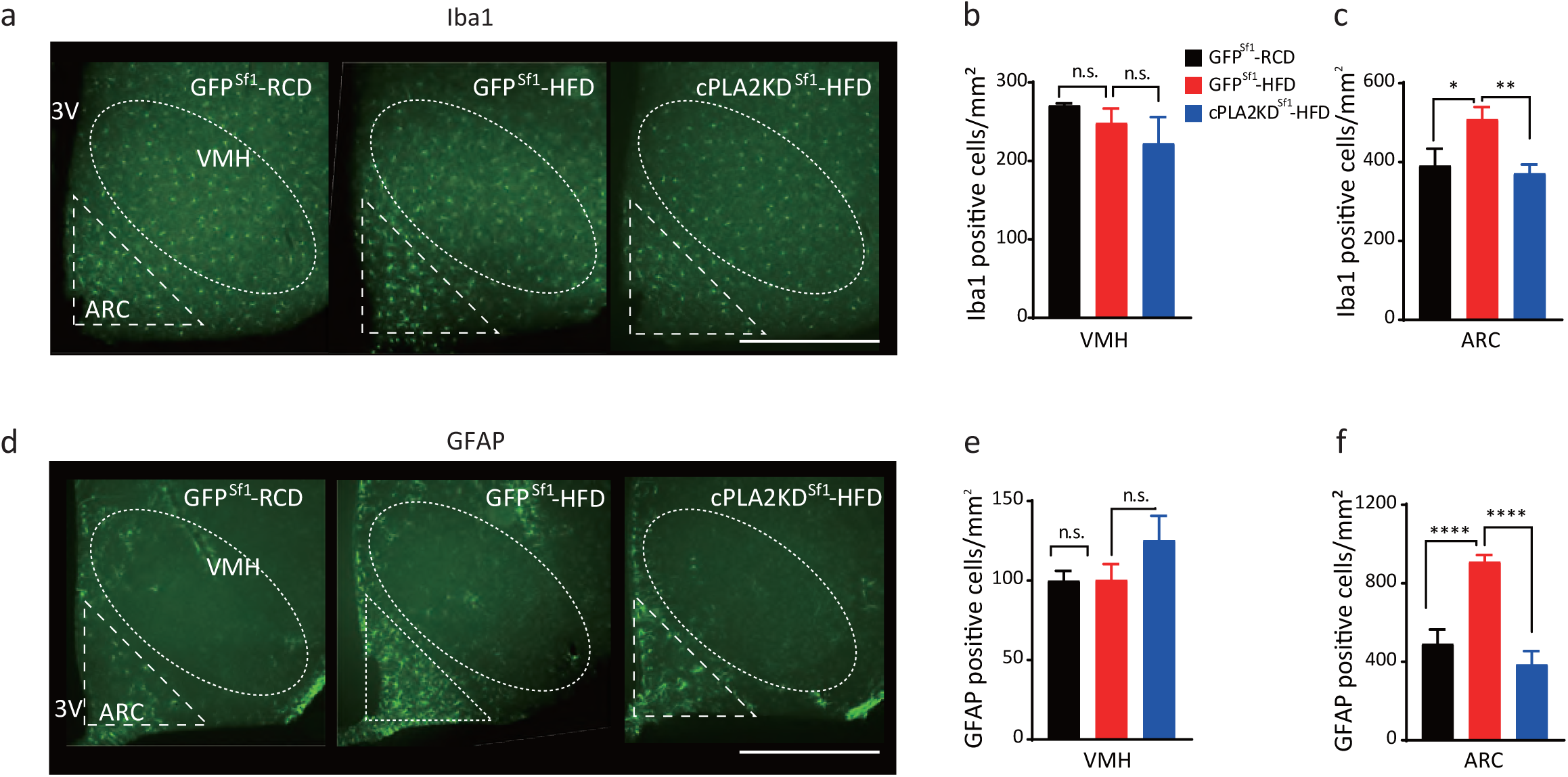
Knockdown of cPLA2 prevents HFD-induced microgliosis and astrogliosis. **a**, Representative micrographs showing immunofluorescent Iba1 staining in the hypothalamus of RCD-fed GFP^Sf1^ mice (GFP^Sf1^-RCD), HFD-fed GFP^Sf1^ mice (GFP^Sf1^-HFD) and HFD-fed cPLA2KD^Sf1^ (cPLA2KD^Sf1^-HFD) mice. Scale bar: 500 μm. **b**,**c**, Quantification of Iba1-positive cells in the VMH (**b**) or ARC (**c**) of GFP^Sf1^-RCD (n=5), GFP^Sf1^-HFD (n=8) and cPLA2KD^Sf1^-HFD (n=5) mice. **d**, Representative micrographs showing immunofluorescent GFAP staining in the hypothalamus of GFP^Sf1^-RCD, GFP^Sf1^-HFD and cPLA2KD^Sf1^-HFD mice. Scale bar: 500 μm. **e**,**f**, Quantification of GFAP-positive cells in the VMH (**e**) or ARC (**f**) of GFP^Sf1^-RCD (n=5), GFP^Sf1^-HFD (n=8) and cPLA2KD^Sf1^-HFD (n=5) mice. All data represent the mean ± SEM; * = p<0.05; ** = p<0.01; *** = p<0.001; **** = p<0.0001.

## Discussion

The roles of hypothalamic phospholipids and eicosanoids in the regulation of energy homeostasis is ill-defined. In this study, we found that the composition of phospholipids in the hypothalamus, especially AA attached phospholipids, are dynamically affected by blood glucose levels. cPLA2 in the VMH plays an important role in AA metabolism to produce prostaglandins and increase insulin sensitivity in muscle during hyperglycemia in RCD-fed mice. cPLA2-mediated phospholipid metabolism also regulates glucose-responsiveness in the dmVMH. HFD feeding, which promotes hyperglycemia, continuously activates cPLA2 and produces prostaglandins, and thus induces inflammation in the hypothalamus and attenuates insulin sensitivity in the liver. Therefore, cPLA2-mediated phospholipid metabolism in the hypothalamus is critical for the physiological and pathological control of systemic glucose homeostasis.

FAs and PUFAs are believed to be transported from the bloodstream to the hypothalamus, and FA metabolism in the hypothalamus changes food intake and energy expenditure^19,27^. However, we observed reductions in AA-containing phospholipids and increases in prostaglandins in the hypothalamus after glucose injection. This suggests that AA is produced from intrinsic membrane phospholipids in the hypothalamus to make eicosanoids during hyperglycemia. The produced prostaglandins play important roles in controlling hyperglycemia because the injection of COX inhibitor impaired glucose tolerance. It has been reported that FA oxidation by carnitine palmitoyltransferase I in the VMH plays important roles in food intake and energy homeostasis^28^. However, our data showed that the cPLA2 in Sf1 neurons has a minor effect on changes in body weight and tissue weight. This suggests that cPLA2 in Sf1 neurons controls glucose metabolism, but not body weight regulation, and it is likely that FAs generated from phospholipids are utilized for prostaglandin production.

AA exists in the sn-2 position of phospholipids, and cPLA2 is the rate-limiting enzyme for catalyzing AA by extracellular stimulation. cPLA2 is activated by an increase in the intracellular calcium concentration and by the phosphorylation of 505-serine residue, which is induced by the MAP kinase pathway^29^. The mechanism that activated cPLA2 in our study remains to be elucidated. However, we found that the glucose-induced activation of the dmVMH is dependent on prostaglandin production by Sf1 neurons. Sf1 neurons exist mainly in the dmVMH and cVMH, and most AA-containing phospholipids were found near the third ventricle in our study. Therefore, the hyperglycemia-induced prostaglandin production occur in the medial part of the hypothalamus and affects neuronal activity in this region probably via changes in ion channel activities^30^. Similarly, prostaglandins regulate glucose-induced insulin secretion (GSIS) from pancreatic beta cells^31^. GSIS is the most studied mechanism of glucose sensing. Thus, it is possible that a similar mechanism for prostaglandins affecting GSIS may be involved in the hypothalamic glucose sensing.

Sf1 neurons are critical for the regulation of whole body energy homeostasis^7,9^. Activation of VMH neurons increases glucose uptake in skeletal muscle and BAT, but not in WAT or other organs^32,33^. Similar results were found in mice with intra-VMH administration of leptin^11,12,34^. Leptin receptors locate in the dmVMH and is required to maintain normal glucose homeostasis^35-37^. In agreement with this, glucose sensing by Sf1 neurons via UCP2 is also critical for systemic glucose metabolism^38^. Therefore, it is plausible that the activation of the dmVMH by glucose injection regulates insulin sensitivity in skeletal muscle through cPLA2-mediated prostaglandin production.

A HFD feeding causes diet-induced-obesity (DIO) and a state of chronic, low-grade inflammation occurs in several tissues, including the hypothalamus^6^. This hypothalamic inflammation is accompanied by an accumulation of microglia, and these changes decrease activities of POMC and AgRP neurons in response to several endocrine signals, such as leptin and insulin^2^. Additionally, a HFD feeding increases astrogliosis in the ARC, paraventrical hypothalamus and dorsomedial hypothalamus, but not the VMH^39^. Consistent with this, our data show that a HFD feeding induces microgliosis and astrogliosis in the ARC but not the VMH, which indicates that the VMH has different inflammatory responses to obesity^39^. After the mice in this study were fed with HFD, all the FAs accumulated in the hypothalamus. Unexpectedly, AA-containing phospholipids decreased because of an increase in hypothalamic cPLA2 activity. Prostaglandins are proinflammatory signals in the brain^40^ and knockdown of cPLA2 in Sf1 neurons attenuates inflammation in the hypothalamus. Our data suggest that the cPLA2-mediated production of prostaglandins in Sf1 neurons enhances inflammatory responses in the whole hypothalamus, including the ARC. It is likely that the long-term production of prostaglandins, which has a physiological role in glucose metabolism in RCD-fed mice, initiates HFD-induced inflammation.

We also found that a HFD feeding abolished the increase in cFos expression induced by glucose in the VMH and ARC, which is consistent with the report that DIO decreases glucose sensing by POMC neurons^41^. In the present study, knockdown of cPLA2 improved the glucose response only in the vlVMH and ARC in HFD-fed mice. However, knockdown of cPLA2 had no effect on the glucose responsiveness of the dmVMH in HFD-fed mice. We also observed that knockdown of cPLA2 impaired glucose sensing in the dmVMH in RCD-fed mice. This indicates that the same attenuation of glucose sensing has already occurred in the dmVMH of HFD-fed mice, thus the dmVMH could not respond to the glucose injection. Our data suggest that inflammation of the hypothalamus contributes to attenuating glucose sensing by the VMH and ARC. POMC and AgRP neurons are reported to regulate hepatic insulin sensitivity, but not muscle glucose metabolism^14,15,42^. Therefore, the improvement of the neuronal activity in the ARC contributes to restoring glucose metabolism by changing insulin sensitivity in the liver.

Aspirin, a COX inhibitor, suppresses insulin sensitivity in healthy human^43–45^, but improves insulin resistance in diabetic patients^46^. Our results were in a good agreement with the human studies. The hypothalamic prostaglandin production may be critical for the effects of aspirin on the whole body insulin sensitivity.

In summary, our study shows that the cPLA2 is fundamental for the function of the hypothalamus in regulating glucose homeostasis. Neuronal cPLA2 is necessary for their own activities in the dmVMH to respond to glucose and control blood glucose levels. However, cPLA2 in the VMH also has the critical role of inducing hypothalamic inflammation during DIO. Therefore, the role of cPLA2-mediated eicosanoid production in the hypothalamus is different between RCD and HFD. Our findings provide novel evidence that cPLA2-mediated phospholipid metabolism in hypothalamic neurons plays an important role in systemic glucose metabolism.

## Methods

### Reagents

All the reagents and resources used in this study are listed in the Supplemental table 1.

### Animals

Sf1-cre mice were purchased from the Jackson Laboratory (STOCK Tg(Nr5a1-cre)7Lowl/J; Bar Harbor, ME). For IMS and assessing the effects of inhibitors, male C57BL6J mice were purchased from Charles River Laboratories Japan. All mice were kept at 22–24 °C with a 12-h light/12-h dark cycle and given *ad libitum* food access. Animal care and experimental procedures were performed with approval from the Animal Care and Use Committee of Hokkaido University.

### Imaging mass spectrometry

Glucose (2 g/kg body weight, Sigma-Aldrich, St. Louis, MO) or saline were injected intraperitoneally (i.p.) and the mice were sacrificed 30 min after injection. Brains were collected and were immediately embedded in 2% sodium carboxymethyl cellulose solution and frozen with liquid nitrogen. The 10-μm brain sections were prepared by cryostat and immediately mounted onto an indium-tin-oxide-coated glass slide (Bruker Daltonics, Bremen, Germany). The sections on the glass slides were immediately dried and stored at −20 °C until imaging mass spectrometry analysis.

Brain slices were sprayed with 9-aminoacridine matrix (10 mg/mL in 70% ethanol, Sigma-Aldrich) and installed into a matrix-assisted laser desorption/ionization (MALDI)-time-of-flight (TOF)/TOF system using ultrafleXtreme (Bruker Daltonics). Brain sections were irradiated by a smart beam (Nd:YAG laser, 355-nm wavelength); with a 25-μm irradiation pitch. The laser had a repetition frequency of 2000 Hz and mass spectra were obtained in the range of *m/z* 200–1200 in negative-ion mode. The *m/z* values from previous reports were used to label each lipid and phospholipid (Supplemental table 2)^47–50^. All ion images were reconstructed with total ion current (TIC) normalization by flexImaging (Bruker Daltonics) and transferred to ImageJ after modifying the grayscale. The areas of the VMH were identified by DAPI staining in the other brain sections and the brightness of the VMH and ARC were calculated as intensity. The intensity ratio in glucose injected mice against saline mice were statistically compared using the Wilcoxon test.

### Quantification of PGs

Glucose (2 g/kg body weight, Sigma-Aldrich) or saline were injected i.p. two times (t = 0 and t = 30 min) and mice were sacrificed at t = 60 min. A HFD (45 kcal% fat, D12451, Research Diet, NJ) was given for 8 weeks. Hypothalamus was collected and immediately frozen in liquid nitrogen. The tissue was homogenized with 500 μl of MeOH:formic acid (100:0.2) containing an internal standard consisting of a mixture of deuterium-labeled PGs using microtip sonication. The samples were submitted to solid phase extraction using an Oasis HLB cartridge (5 mg; Waters, Milford, MA) according to the method of Kita et al^51^. Briefly, samples were diluted with water:formic acid (100:0.03) to give a final MeOH concentration of ∼ 20% by volume, applied to preconditioned cartridges, and washed serially with water:formic acid (100:0.03), water:ethanol:formic acid (90:10:0.03), and petroleum ether. Samples were eluted with 200 μl of MeOH:formic acid (100:0.2). The filtrate was concentrated with a vacuum concentrator (SpeedVac, Thermo Fisher Scientific, Waltham, MA). The concentrated filtrate was dissolved in 50 μL of methanol and used for liquid chromatography/mass spectrometry (LC-MS).

Hypothalamic PGs were quantified by modified LC-MS^52^. Briefly, a triple-quadrupole mass spectrometer equipped with an electrospray ionization (ESI) ion source (LCMS-8060; Shimadzu Corporation, Kyoto, Kyoto, Japan) was used in the positive and negative-ESI and multiple reaction monitoring modes.

### Stereotaxic surgeries and AAV injection

Male C57BL/6J mice were anesthetized with mixture of ketamine (100 mg/kg) and xylazine (10 mg/kg) and were put on a stereotaxic instrument (Narishige, Tokyo, Japan). Mice were implanted with cannulae for intracerebroventricular (i.c.v.) or intra-hypothalamic injection. The i.c.v. cannulae were implanted in the lateral ventricle in an anterior–posterior (AP) direction: −0.3 (0.3 mm posterior to the bregma), lateral (L): 1.0 (1.0 mm lateral to the bregma), dorsal–ventrol (DV): −2.5 (2.5 mm below the bregma on the surface of the skull). The double-cannulae for intrahypothalamic injection had a gap of 0.8 mm between the two cannulae and were implanted following the coordinates of the AP: −1.4, L: ± 0.4, DV: −5.6. Cannulae were secured on the skulls with cyanoacrylic glue and the exposed skulls were covered with dental cement. To knock down expression of *pla2g4a* in the VMH, 6- to 8-week-old Sf1-cre mice were injected in each side of the VMH with ∼0.5 µL AAV8-DIO-shRNA (Vigene Biosciences, Rockville, MD) against *mpla2g4* bilaterally using the following coordinates: AP: −1.4, L: ± 0.4, DV: −5.7. Open wounds were sutured after viral injection. Mice were allowed to recover for 5–7 days before experiments were started.

### Glucose and insulin tolerance tests

A glucose tolerance test was performed on *ad libitum* fed or fasted mice. The *ad libitum* fed mice were used for assessing the effects of inhibitors. The fasted mice were used for assessing the phenotype of mice with knockdown of *pla2g4a* in Sf1-neurons (cPLA2KD^Sf1^). To assess the effects of inhibitors, methyl arachidonyl fluorophosphonate (MAFP; 20 μM, 300 nL in each side), indomethacin (140 μM, 300 nL in each side), or vehicle were injected into both sides of the hypothalamus through a double-cannula. Glucose solution was then injected i.p. (2 g/kg) 30 min after intrahypothalamic injection. To assess the phenotype of cPLA2KD^Sf1^ mice, animals were fasted for 16 hours and injected with glucose (2 g/kg) i.p.. Blood glucose levels were measured by a handheld glucose meter (Nipro Free style, Nipro, Osaka, Japan) before injecting inhibitors (−30 min) or glucose (0 min), and measured at 15, 30, 60 and 120 min after glucose injection.

An insulin tolerance test was performed in *ad libitum* fed mice. The mice were i.p. injected with 0.5 U/kg insulin (Novo Nordisk, Bagsværd, Denmark). Blood glucose was measured before injecting inhibitors (−30 min) and glucose (0 min), and measured 15, 30, 60 and 120 min after glucose injection.

### Serum insulin measurement

Mice were injected with inhibitors or vehicle into the hypothalamus using the same protocol as above. Blood from the tails was taken 30 min after intrahypothalamic injection. Then, glucose (2 g/kg) was i.p. injected and blood was taken at 15 and 30 min after glucose injection. The serums were collected after centrifuging for 10 min at 1000 ×g and maintained at −80°C until insulin was measured. The insulin concentration was measured with a Mouse Insulin ELISA KIT (FUJIFILM Wako, Osaka, Japan) and all procedures were followed by the protocols provided in the kit.

### Real time PCR

Total RNA was extracted from the whole hypothalamus using Trizol solution (Invitrogen). *Pla2g4, Rbfox3* and *Actb* mRNA levels in the hypothalamus were measured by real-time TaqMan PCR. A high capacity cDNA reverse transcription kit (Thermo Fisher Scientific) was used for the reverse transcription. Real-time PCR (LightCycler 480; Roche) was performed with diluted cDNAs in a 20-μl reaction volume in triplicate.

### Immunohistochemistry

Ad libitum fed mice were i.p. injected with either saline or glucose (3 g/kg) and perfused with heparinized saline followed by 4% paraformaldehyde (PFA) transcardially at 30 min after injection. Inhibitors were i.c.v. injected 30 min before glucose injection. Brain sections (50 μm each) containing the whole VMH were collected. Floating sections were incubated with rabbit-anti-cFos antibody (1:200, Santa Cruz Biotechnology, Denton, TX) or rabbit-anti-GFP antibody (1:1000, Frontier Institute, Hokkaido, Japan) in staining solution (0.1 M phosphate buffer (PB) containing 4% normal guinea pig serum, 0.1% glycine, and 0.2% Triton X-100) overnight at room temperature. To assess the cell population of astrocytes and microglia, sections were incubated with rabbit-anti-Iba1 antibody (1:3000, FUJIFILM Wako) or rabbit-anti-GFAP antibody (1:3000, Sigma-Aldrich) in staining solution overnight at room temperature. After rinsing with PB, sections were incubated in secondary antibody (1:500, Alexa Fluor 647 or 488 Goat Anti-Rabbit (IgG) secondary antibody, Cell Signaling Technologies, Danvers, MA) for 2 h at room temperature. The stained sections were washed with PB three times and mounted on glass slides with vectashield (Vector Laboratories, Burlingame, CA).

### Assessment of cytosolic- or secretory-phospholipase-A2 activity

Mice fed *ad libitum* were i.p. injected with either glucose (2 g/kg) or saline. Mouse hypothalami were collected 30 min after injection and stored at −80 °C until use. Tissues were homogenized and centrifuged at 10,000 ×g for 15 min at 4 °C and supernatants were collected. Activity of cytosolic- or secretory-phospholipase-A2 were measured following procedures described in the kit manuals (Abcam, Cambridge, UK).

### Implantation of artery and vein catheter for clamp studies

Mice were anesthetized with pre-mixed ketamine (100 mg/kg) and xylazine (10 mg/kg). Polyethylene catheters were implanted into right carotid arteries and jugular veins. The tubes entered subcutaneously and protruded from the neck skin. Mice were allowed to recover for 3 to 5 days and tubes were flushed with heparinized saline each day.

### Hyperinsulinemic–euglycemic clamp and measurement of 2-[14C] deoxy-D-glucose (2DG) uptake

The hyperinsulinemic–euglycemic clamp protocol was followed as described in previous papers^37,53^. The mice were fasted for 4 h and experiments were initiated in a free moving condition.

A 115-min clamp period (t = 0–115 min) was following a 90-min basal period (t = −90 to 0 min). A bolus of [3-^3^H] glucose (5 mCi;) was injected through the jugular vein at the beginning of the basal period (t = −90 min) and tracer was infused at a rate of 0.05 mCi for 90 min. Blood samples were collected at t = −15 and −5 min to measure the rate of appearance (Ra). The clamp period was initiated with continuous infusion of insulin (2.5 mU/kg/min). During the clamp period, blood was collected and blood glucose levels were measured from arterial blood every 5–10 min. Cold glucose was infused at a variable rate via the jugular vein catheter to maintain a blood glucose level at 110–130 mg/dL. Erythrocytes in withdrawn blood were suspended in sterile saline and returned to each animal.

To assess 2DG uptake, 2-[^14^C] DG (10 mCi) was infused at t = 70 min and blood samples were collected at t = 75, 85, 95, 105, and 115 min. After collecting the blood sample at t = 115 min, mice were euthanatized, and small pieces of tissue samples from the soleus, Gastro-R (red portion of gastrocnemius), Gastro-W (white portion of gastrocnemius), BAT, heart, spleen, EWAT (epididymal white adipose tissue), brain (cortex), and liver were rapidly collected. The rate of disappearance (Rd), which reflects whole-body glucose utilization, rate of appearance (Ra), which mainly reflects endogenous glucose production (EGP), and the rates of whole-body glycolysis and glycogen synthesis were determined as described previously^53^.

### Statistical Analysis

Two-way or one-way ANOVA were used to determine the effect of inhibitors or knockdown of cPLA2 with the Prism 8 software (GraphPad). For repeated-measures analysis, ANOVA was used when values over different times were analyzed, followed by the Bonferroni and Sidak multiple comparisons tests. When only two groups were analyzed, statistical significance was determined by the unpaired Student’s t test. A value of p<0.05 was considered statistically significant. All data are shown as mean ± SEM.

### Data availability

The data that support the findings of this study are available from the corresponding author upon reasonable request.

## Acknowledgments

This work was supported by Leading Initiative for Excellent Young Researchers (from MEXT); a Grant-in-Aid for Young Scientists (A) (Grant Number JP17H05059), a Grant-in-Aid for Scientific Research (B) (Grant Number JP18H02857); Japanese Initiative for Progress of Research on Infectious Diseases for Global Epidemics (JP17fm0208011h0001, JP18fm0208011h0002, JP19fm0208011h0003); the Takeda Science Foundation; the Uehara Memorial Foundation; Astellas Foundation for Research on Metabolic Disorders; Suzuken Memorial Foundation; Program for supporting introduction of the new sharing system (JPMXS0420100617, JPMXS0420100618, JPMXS0420100619) and National Institutes of Health (RO1 DK107293). Infrastructure of LC-MS was supported by JST ERATO Suematsu Gas Biology Project. M.S. is the lead until March 2015.

## Author Contributions

C.T. conceived this study and designed the experiments. M.L. performed most of the experiments and C.T. supervised the entire study. H.M. performed the study on astrocytes. Y.S. performed LC-MS measurements. T.H. performed imaging mass spectrometry. I.Y., D.I. performed GTT, ITT and AAV injections. M.S. N.I., K.K., S.D. assisted in preparing the manuscript.

## Competing interests

The authors declare no competing interests.

## Materials & Correspondence

Further information and requests for resources and reagents should be directed to and will be fulfilled by the Lead Contact, Chitoku Toda.

## Supplemental Information

**Supplemental Figure 1**

Relative amounts of hypothalamic eicosanoids mediated by lipoxygenase (**a**) or cytochrome P450 (**b**) after the injection of glucose compared with saline injected mice. n=3 in each experimental group. Data represent the mean fold change in color.

**Supplemental Figure 2**

Hypothalamic PLC-mediated-pathway does not affect systemic glucose metabolism. (**a**) Glucose tolerance test (GTT) (0–120 min) after intra-hypothalamic injection (−30 min) of Xestospondin, an IP3 receptor antagonist, (n=7) or vehicle (n=7). (**b**) GTT (0–120 min) after intra-hypothalamic injection (−30 min) of U73122, a phospholipase C (PLC) inhibitor, (n=7) or vehicle (n=7). All data represent the mean ± SEM

**Supplemental Figure 3**

**a**, Construct of AAV8-DIO (CreOn)-shRNA against *mpla2g4*, containing DIO (Double-floxed Inverted Open reading frame) to express shRNA Cre-dependently. **b**, Representative micrographs showing virus infected (tdTomato) and shRNA expressing (GFP) Sf1-neurons. Scale bar: 500 μm. **c**, Expression of *pla2g4a* mRNA in the whole hypothalamus injected with AAV8-DIO-shRNA against *mpla2g4* (cPLA2KD^Sf1^; n=3) compared with control mice (GFP^Sf1^; n=3). **d**, Body weight and tissue weight in cPLA2KD^Sf1^ mice (n=5) and GFP^Sf1^ mice (n=5). (IWAT: inguinal white adipose tissue. EWAT: epididymal white adipose tissue. mesentWAT: mesenteric white adipose tissue. BAT: brown adipose tissue.) ll data represent the mean ± SEM

**Supplemental Figure 4**

Knockdown of astrocytic cPLA2 in hypothalamus (cPLA2KD^GFAP^) did not change body weight and glucose metabolism. **a**, Construct of AAV8-GFAP-Cre-mCherry and AAV8-DIO-shRNA against *mpla2g4* and representative micrographs showing virus infected (tdTomato) and shRNA expressing (GFP) astrocytes in the ARC. Scale bar: 25 μm. **b**, Relative expression of cPLA2 in the hypothalamus of cPLA2KD^GFAP^ and control mice. n=3 in each experimental group. **c**, Glucose tolerance test in cPLA2KD^GFAP^ mice (n=10) and GFP^GFAP^ mice (n=9). **d**, Insulin tolerance test in cPLA2KD^GFAP^ mice (n=10) and GFP^GFAP^ mice (n=9). **e**, Body weight change in cPLA2KD^GFAP^ mice (n=10) and GFP^GFAP^ mice (n=9) after viral injection. All data represent the mean ± SEM.

**Supplemental Figure 5**

Relative amounts of hypothalamic eicosanoids mediated by lipoxygenase (**a**) or cytochrome P450 (**b**) in RCD or HFD fed mice. n=3 each experimental group. Data represent the mean fold change in color.

**Supplemental Figure 6**

Body weight and tissue weight in cPLA2KD^Sf1^ mice (n=5) and GFP^Sf1^ mice (n=5) after 8 weeks of HFD feeding. (IWAT: inguinal white adipose tissue. EWAT: epididymal white adipose tissue. mesentWAT: mesenteric white adipose tissue. BAT: brown adipose tissue.)

**Supplemental table 1**

List of reagents and resources

**Supplemental table 2**

Assignment of lipid molecular species by IMS negative ion mode

## Graphical abstract

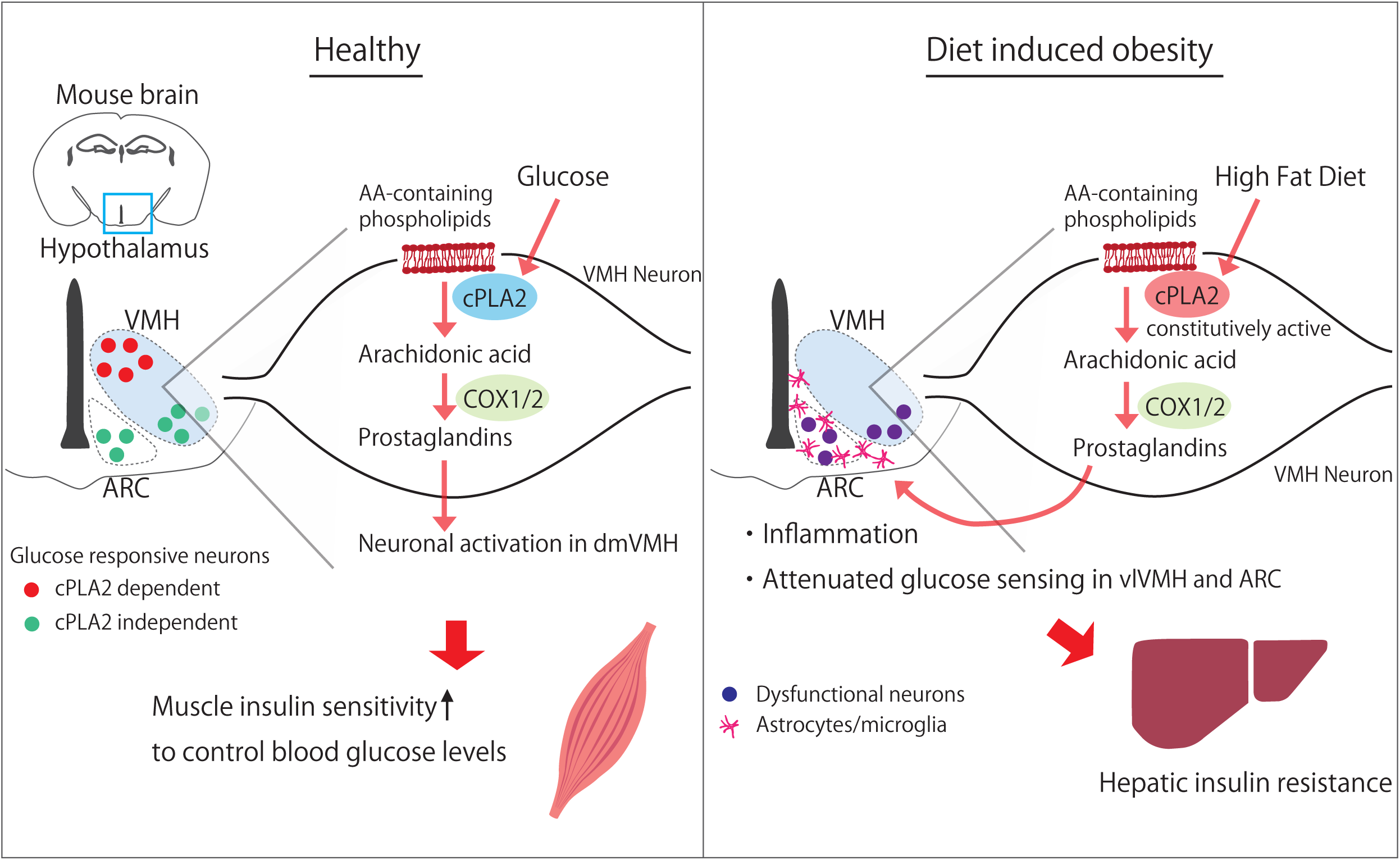

## Notes

### Competing Interest Statement

The authors have declared no competing interest.

## References

1. Pozo, M. & Claret, M. Hypothalamic Control of Systemic Glucose Homeostasis: The Pancreas Connection. Trends in Endocrinology & Metabolism 29, 581–594 (2018).

2. Ruud, J., Steculorum, S. M. & Brüning, J. C. Neuronal control of peripheral insulin sensitivity and glucose metabolism. Nature Communications 8, 15259 (2017).

3. Garfield, A. S. et al. A Parabrachial-Hypothalamic Cholecystokinin Neurocircuit Controls Counterregulatory Responses to Hypoglycemia. Cell Metabolism 20, 1030–1037 (2014).

4. Meek, T. H. et al. Functional identification of a neurocircuit regulating blood glucose. PNAS 113, E2073–E2082 (2016).

5. Myers, M. G. & Olson, D. P. Central nervous system control of metabolism. Nature 491, 357–363 (2012).

6. Cai, D. & Khor, S. “Hypothalamic Microinflammation” Paradigm in Aging and Metabolic Diseases. Cell Metabolism 30, 19–35 (2019).

7. Shimazu, T. & Minokoshi, Y. Systemic Glucoregulation by Glucose-Sensing Neurons in the Ventromedial Hypothalamic Nucleus (VMH). Journal of the Endocrine Society 1, 449–459 (2017).

8. Stanley, S. A. et al. Bidirectional electromagnetic control of the hypothalamus regulates feeding and metabolism. Nature 531, 647–650 (2016).

9. Coutinho, E. A. et al. Activation of SF1 Neurons in the Ventromedial Hypothalamus by DREADD Technology Increases Insulin Sensitivity in Peripheral Tissues. Diabetes 66, 2372–2386 (2017).

10. Minokoshi, Y., Haque, M. S. & Shimazu, T. Microinjection of leptin into the ventromedial hypothalamus increases glucose uptake in peripheral tissues in rats. Diabetes 48, 287–291 (1999).

11. Toda, C. et al. Distinct effects of leptin and a melanocortin receptor agonist injected into medial hypothalamic nuclei on glucose uptake in peripheral tissues. Diabetes 58, 2757–2765 (2009).

12. Toda, C. et al. Extracellular Signal–Regulated Kinase in the Ventromedial Hypothalamus Mediates Leptin-Induced Glucose Uptake in Red-Type Skeletal Muscle. Diabetes 62, 2295–2307 (2013).

13. Roh, E. & Kim, M.-S. Brain Regulation of Energy Metabolism. Endocrinol Metab (Seoul) 31, 519–524 (2016).

14. Dodd, G. T. et al. Insulin regulates POMC neuronal plasticity to control glucose metabolism. eLife 7, e38704 (2018).

15. Könner, A. C. et al. Insulin Action in AgRP-Expressing Neurons Is Required for Suppression of Hepatic Glucose Production. Cell Metabolism 5, 438–449 (2007).

16. Routh, V. H., Hao, L., Santiago, A. M., Sheng, Z. & Zhou, C. Hypothalamic glucose sensing: making ends meet. Frontiers in Systems Neuroscience 8, 236 (2014).

17. Loftus, T. M. et al. Reduced Food Intake and Body Weight in Mice Treated with Fatty Acid Synthase Inhibitors. Science 288, 2379–2381 (2000).

18. Pocai, A., Obici, S., Schwartz, G. J. & Rossetti, L. A brain-liver circuit regulates glucose homeostasis. Cell Metabolism 1, 53–61 (2005).

19. Bazinet, R. P. & Layé, S. Polyunsaturated fatty acids and their metabolites in brain function and disease. Nat. Rev. Neurosci. 15, 771–785 (2014).

20. Ghosh, M., Tucker, D. E., Burchett, S. A. & Leslie, C. C. Properties of the Group IV phospholipase A2 family. Progress in Lipid Research 45, 487–510 (2006).

21. Nonogaki, K. et al. Dissociation of hyperthermic and hyperglycemic effects of central prostaglandin F2α. Prostaglandins 41, 451–462 (1991).

22. Migrenne, S. et al. Fatty Acid Signaling in the Hypothalamus and the Neural Control of Insulin Secretion. Diabetes 55, S139–S144 (2006).

23. Obici, S. et al. Central Administration of Oleic Acid Inhibits Glucose Production and Food Intake. Diabetes 51, 271–275 (2002).

24. Farooqui, A. A., Yang, H.-C., Rosenberger, T. A. & Horrocks, L. A. Phospholipase A2 and Its Role in Brain Tissue. Journal of Neurochemistry 69, 889–901 (1997).

25. Macedo, F., dos Santos, L. S., Glezer, I. & da Cunha, F. M. Brain Innate Immune Response in Diet-Induced Obesity as a Paradigm for Metabolic Influence on Inflammatory Signaling. Front Neurosci 13, 342 (2019).

26. Posey, K. A. et al. Hypothalamic proinflammatory lipid accumulation, inflammation, and insulin resistance in rats fed a high-fat diet. American Journal of Physiology-Endocrinology and Metabolism 296, E1003–E1012 (2009).

27. Rapoport, S. I., Chang, M. C. & Spector, A. A. Delivery and turnover of plasma-derived essential PUFAs in mammalian brain. J. Lipid Res. 42, 678–685 (2001).

28. Bruce, K. D., Zsombok, A. & Eckel, R. H. Lipid Processing in the Brain: A Key Regulator of Systemic Metabolism. Front Endocrinol (Lausanne) 8, 60 (2017).

29. Lin, L. L. et al. cPLA2 is phosphorylated and activated by MAP kinase. Cell 72, 269–278 (1993).

30. Jang, Y., Kim, M. & Hwang, S. W. Molecular mechanisms underlying the actions of arachidonic acid-derived prostaglandins on peripheral nociception. J Neuroinflammation 17, 30 (2020).

31. Carboneau, B. A., Breyer, R. M. & Gannon, M. Regulation of pancreatic β-cell function and mass dynamics by prostaglandin signaling. J Cell Commun Signal 11, 105–116 (2017).

32. Shimazu, T., Sudo, M., Minokoshi, Y. & Takahashi, A. Role of the hypothalamus in insulin-independent glucose uptake in peripheral tissues. Brain Res. Bull. 27, 501–504 (1991).

33. Sudo, M., Minokoshi, Y. & Shimazu, T. Ventromedial hypothalamic stimulation enhances peripheral glucose uptake in anesthetized rats. Am. J. Physiol. 261, E298–303 (1991).

34. Kamohara, S., Burcelin, R., Halaas, J. L., Friedman, J. M. & Charron, M. J. Acute stimulation of glucose metabolism in mice by leptin treatment. Nature 389, 374–377 (1997).

35. Dhillon, H. et al. Leptin Directly Activates SF1 Neurons in the VMH, and This Action by Leptin Is Required for Normal Body-Weight Homeostasis. Neuron 49, 191–203 (2006).

36. Sohn, J.-W. et al. Leptin and insulin engage specific PI3K subunits in hypothalamic SF1 neurons. Molecular Metabolism 5, 669–679 (2016).

37. Zhang, R. et al. Selective Inactivation of Socs3 in SF1 Neurons Improves Glucose Homeostasis without Affecting Body Weight. Endocrinology 149, 5654–5661 (2008).

38. Toda, C. et al. UCP2 Regulates Mitochondrial Fission and Ventromedial Nucleus Control of Glucose Responsiveness. Cell 164, 872–883 (2016).

39. Buckman, L. B., Thompson, M. M., Moreno, H. N. & Ellacott, K. L. J. Regional astrogliosis in the mouse hypothalamus in response to obesity. Journal of Comparative Neurology 521, 1322–1333 (2013).

40. Palumbo, S. & Bosetti, F. Alterations of brain eicosanoid synthetic pathway in multiple sclerosis and in animal models of demyelination: Role of cyclooxygenase-2. Prostaglandins, Leukotrienes and Essential Fatty Acids 89, 273–278 (2013).

41. Parton, L. E. et al. Glucose sensing by POMC neurons regulates glucose homeostasis and is impaired in obesity. Nature 449, 228–232 (2007).

42. Berglund, E. D. et al. Direct leptin action on POMC neurons regulates glucose homeostasis and hepatic insulin sensitivity in mice. J Clin Invest 122, 1000–1009 (2012).

43. Bratusch-Marrain, P. R., Vierhapper, H., Komjati, M. & Waldhäusl, W. K. Acetyl-salicylic acid impairs insulin-mediated glucose utilization and reduces insulin clearance in healthy and non-insulin-dependent diabetic man. Diabetologia 28, 671–676 (1985).

44. Giugliano, D., Sacca, L., Scognamiglio, G., Ungaro, B. & Torella, R. Influence of acetylsalicylic acid on glucose turnover in normal man. Diabete Metab 8, 279–282 (1982).

45. Newman, W. P. & Brodows, R. G. Aspirin causes tissue insensitivity to insulin in normal man. J. Clin. Endocrinol. Metab. 57, 1102–1106 (1983).

46. Hundal, R. S. et al. Mechanism by which high-dose aspirin improves glucose metabolism in type 2 diabetes. J. Clin. Invest. 109, 1321–1326 (2002).

47. Cerruti, C. D., Benabdellah, F., Laprévote, O., Touboul, D. & Brunelle, A. MALDI Imaging and Structural Analysis of Rat Brain Lipid Negative Ions with 9-Aminoacridine Matrix. Anal. Chem. 84, 2164–2171 (2012).

48. Goto, T. et al. The expression profile of phosphatidylinositol in high spatial resolution imaging mass spectrometry as a potential biomarker for prostate cancer. PLoS ONE 9, e90242 (2014).

49. Rocha, B. et al. Characterization of lipidic markers of chondrogenic differentiation using mass spectrometry imaging. Proteomics 15, 702–713 (2015).

50. Fülöp, A. et al. 4-Phenyl-α-cyanocinnamic acid amide: screening for a negative ion matrix for MALDI-MS imaging of multiple lipid classes. Anal. Chem. 85, 9156–9163 (2013).

51. Kita, Y., Takahashi, T., Uozumi, N. & Shimizu, T. A multiplex quantitation method for eicosanoids and platelet-activating factor using column-switching reversed-phase liquid chromatography–tandem mass spectrometry. Analytical Biochemistry 342, 134–143 (2005).

52. Yamada, M. et al. A comprehensive quantification method for eicosanoids and related compounds by using liquid chromatography/mass spectrometry with high speed continuous ionization polarity switching. Journal of Chromatography B 995–996, 74–84 (2015).

53. Ayala, J. E., Bracy, D. P., McGuinness, O. P. & Wasserman, D. H. Considerations in the Design of Hyperinsulinemic-Euglycemic Clamps in the Conscious Mouse. Diabetes 55, 390–397 (2006).

