## Supplemental figures and tables for "Prostaglandin in the ventromedial hypothalamus regulates peripheral glucose metabolism"

supplementary figure1

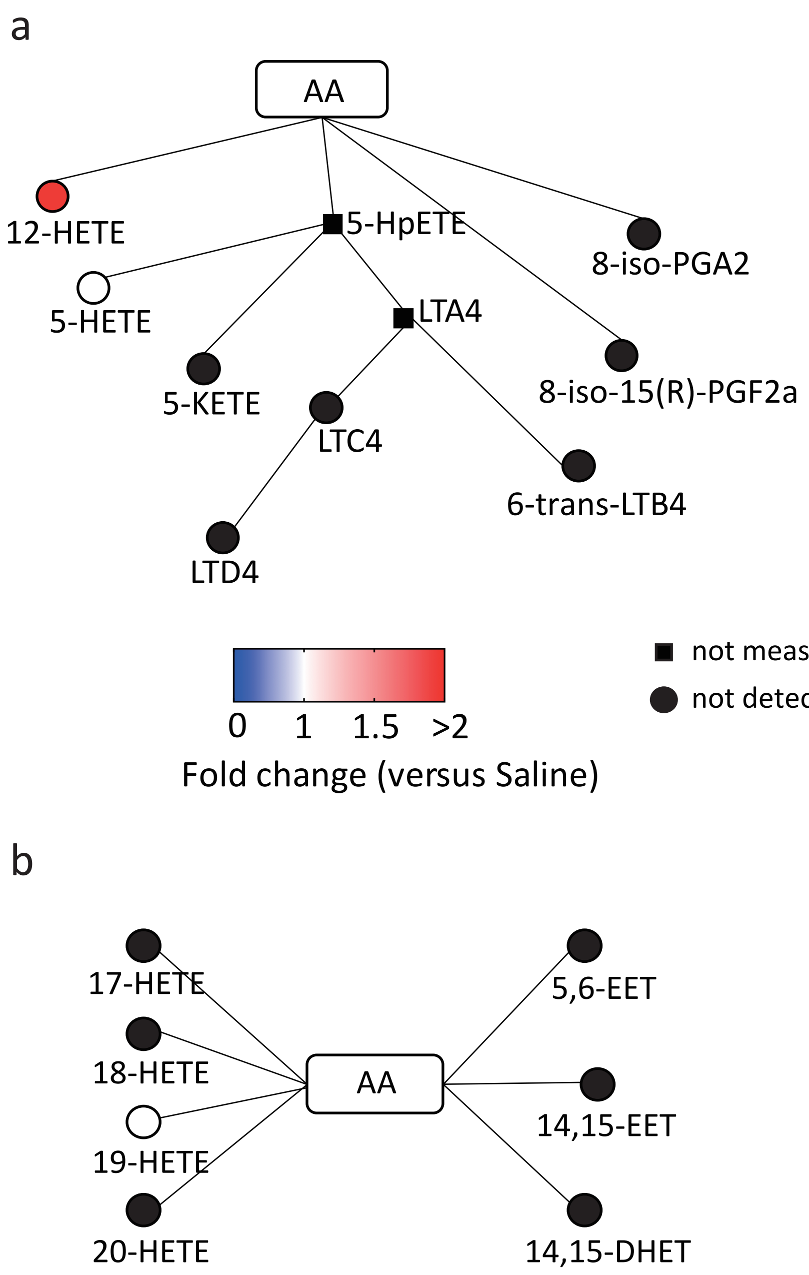


Supplemental Figure 1
Relative amounts of hypothalamic eicosanoids mediated by lipoxygenase (**a**) or cytochrome P450 (**b**) after the injection of glucose compared with saline injected mice. n=5 in each experimental group. Data represent the mean fold change in color.

supplementary figure2

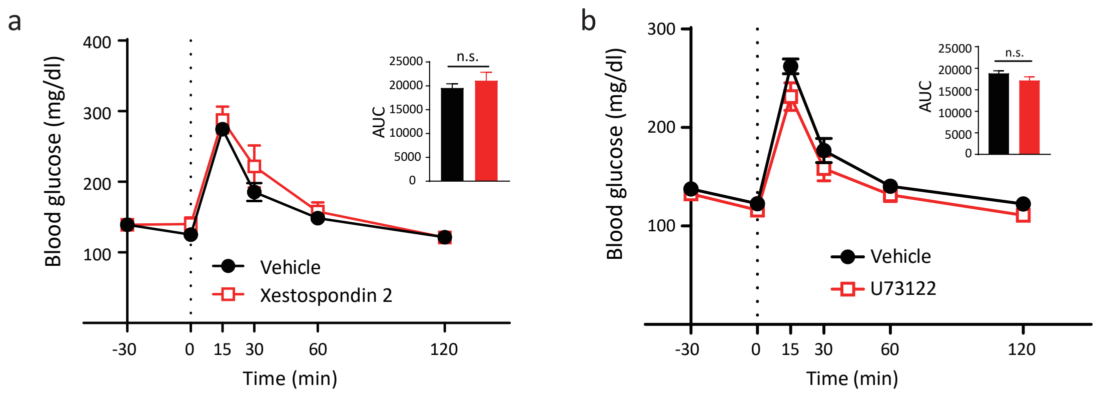


Supplemental Figure 2

Hypothalamic PLC mediated-pathway does not affect systemic glucose metabolism. **a,** Glucose tolerance test (GTT) (0–120 min) after intra-hypothalamic injection (-30 min) of Xestospondin, a IP3 receptor antagonist, (n=7) or vehicle (n=7). **b,** GTT (0–120 min) after intra-hypothalamic injection (-30 min) of U73122, a phospholipase C (PLC) inhibitor, (n=7) or vehicle (n=7). All data represent the mean ± SEM

supplementary figure3

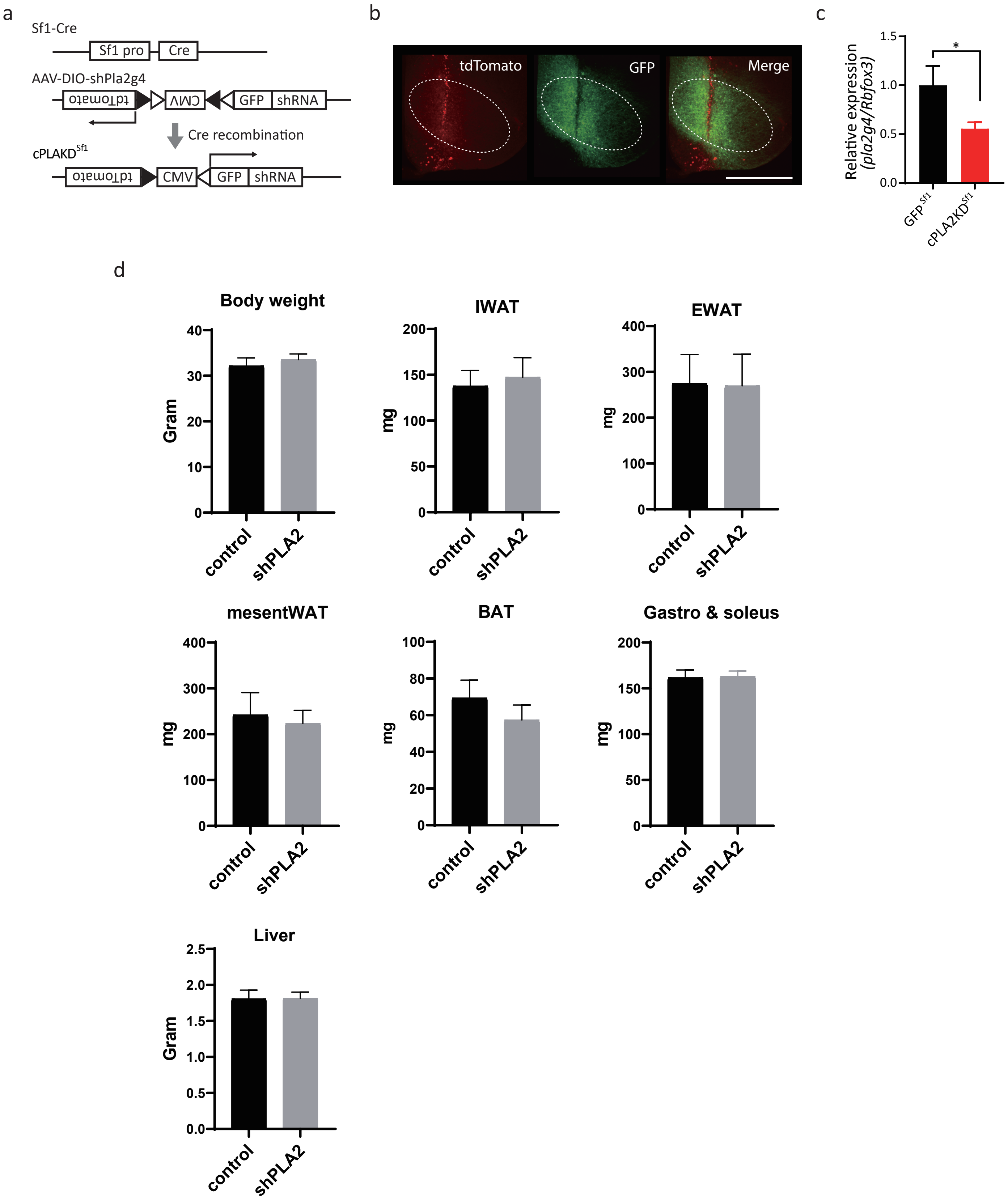


Supplemental Figure 3

**a,** Construct of AAV8-DIO (CreOn)-shRNA against mpla2g4, containing DIO (Double-floxed Inverted Open reading frame) to express shRNA Cre-dependently. **b,** Representative micrographs showing virus infected (tdTomato) and shRNA expressing (GFP) Sf1-neurons. Scale bar: 500 µm. **c,** Expression of pla2g4a mRNA in the whole hypothalamus injected with AAV8-DIO-shRNA against mpla2g4 (cPLA2KD$^{Sf1}$; n=3) compared with control mice (GFP$^{Sf1}$; n=3). **d,** Body weight and tissue weight in cPLA2KD$^{Sf1}$ mice (n=5) and GFP$^{Sf1}$ mice (n=5). (IWAT: inguinal white adipose tissue. EWAT: epididymal white adipose tissue. mesentWAT: mesenteric white adipose tissue. BAT: brown adipose tissue.)
All data represent the mean ± SEM

supplementary figure 4

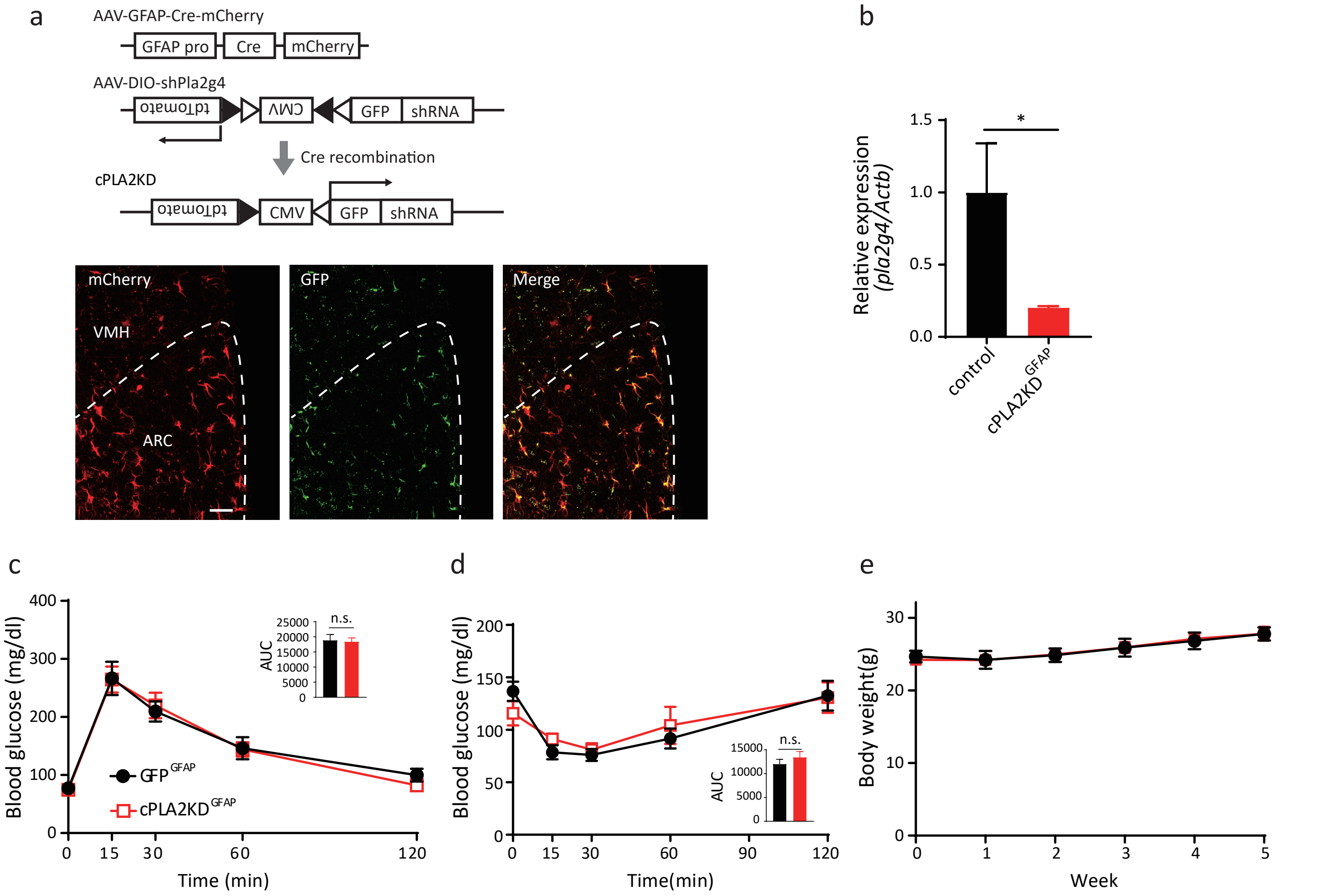


Supplemental Figure 4
Knockdown of astrocytic cPLA2 in hypothalamus (cPLA2KD$^{GFAP}$) did not change body weight and glucose metabolism. **a,** Construct of AAV8-GFAP-Cre-mCherry and AAV8-DIO-shRNA against mpla2g4 and representative micrographs showing virus infected (tdTomato) and shRNA expressing (GFP) astrocytes in the ARC. Scale bar: 25 μm. **b,** Relative expression of cPLA2 in the hypothalamus of cPLA2KD$^{GFAP}$ and control mice. n=3 in each experimental group. **c,** Glucose tolerance test in cPLA2KD$^{GFAP}$ mice (n=10) and GFP$^{GFAP}$ mice (n=9). **d,** Insulin tolerance test in cPLA2KD$^{GFAP}$ mice (n=10) and GFP$^{GFAP}$ mice (n=9). **e,** Body weight change in cPLA2KD$^{GFAP}$ mice (n=10) and GFP$^{GFAP}$ mice (n=9) after viral injection. All data represent the mean ± SEM.

supplementary figure 5

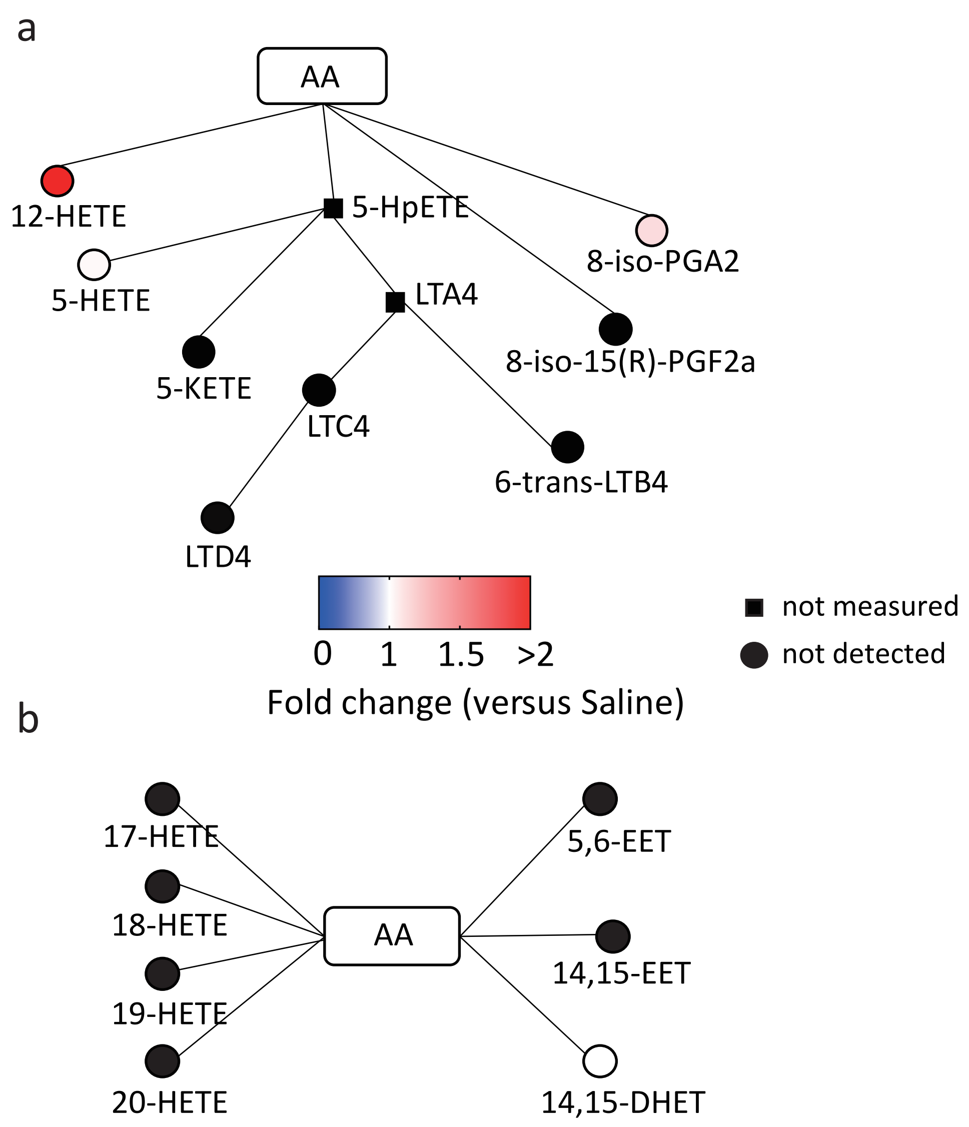


Supplemental Figure 5
Relative amounts of hypothalamic eicosanoids mediated by lipoxygenase (**a**) or cytochrome P450 (**b**) in RCD or HFD fed mice. n=3 each experimental group. Data represent the mean fold change in color.

Supplemental Figure 6

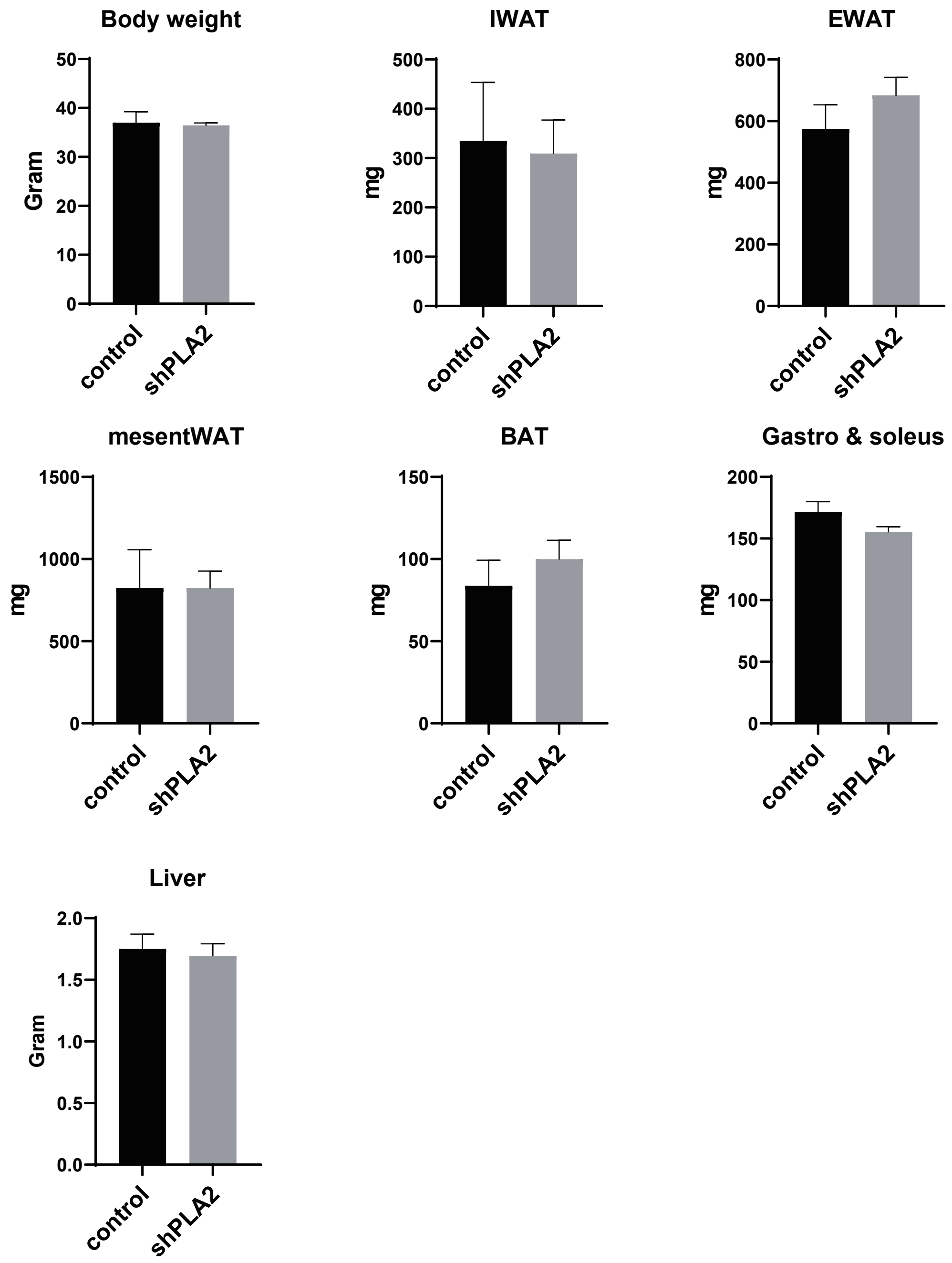


Supplemental Figure 6
Body weight and tissue weight in cPLA2KD$^{Sf1}$ mice (n=5) and GFP$^{Sf1}$ mice (n=5) after 8 weeks of a HFD feeding. (IWAT: inguinal white adipose tissue. EWAT: epididymal white adipose tissue. mesentWAT: mesenteric white adipose tissue. BAT: brown adipose tissue.)

supplemental table 1

List of reagents and resources

| REAGENT or RESOURCE | SOURCE | IDENTIFIER |
|---|---|---|
| Antibodies | | |
| Rabbit-anti-cFos | Santa Cruz | CAT#SC-52 |
| Rabbit-anti-Iba1 | FUJIFILM Wako | CAT#LKN5648 |
| Rabbit-anti-GFAP | Sigma-Aldrich | CAT#HPA050030 |
| Rabbit-anti-GFP | Frontier institute | AB_2571573 |
| Anti-rabbit IgG (H+L), F(ab')2 Fragment (Alexa Fluor 647 Conjugate) | Cell Signaling Technologies | 4414S |
| Anti-rabbit IgG (H+L), F(ab')2 Fragment (Alexa Fluor® 488 Conjugate) | Cell Signaling Technologies | 4412S |
| Bacterial and Virus Strains | | |
| AAV8-DIO (Cre-On)-shRNA against *mpla2g4* | Vigene | |
| AAV8-GFAP-mcherry-Cre | UNC Vector Core | Lot# AV5056C |
| Chemicals, Peptides, and Recombinant Proteins | | |
| 9-Aminoacridine hemihydrate | Thermo Fisher Scientific | 134410010 |
| Glucose | FUJIFILM Wako | 049-31165 |
| Novolin R 100 IU | Novo Nordisk | N/A |
| Glucose-D-[3-$^{3}$H] | Muromachi Kikai | ART0124 |
| 2-Deoxy-D-glucose $^{14}$C(U) | Muromachi Kikai | ARC0112A |
| Methyl arachidonyl fluorophosphonate (MAFP) | Sigma-Aldrich | M2939 |
| Indomethacin (Indo) | Sigma-Aldrich | I7378 |
| Ethanol | FUJIFILM Wako | 056-03341 |
| Dimethyl sulfoxide | Nacalai Tesque | 13407 |
| TRIzol™ reagent | Thermo Fisher Scientific | 15596026 |
| Critical Commercial Assays | | |
| Mouse Insulin ELISA KIT | FUJIFILM Wako | 633-23919 |
| Cytosolic Phospholipase A2 Assay Kit | Abcam | Ab133090 |
| Secretory-phospholipase-A2 Assay Kit | Abcam | Ab133089 |
| TaqMan™ Gene Expression Master Mix | Thermo Fisher Scientific | 4369016 |
| M-MLV Reverse Transcriptase | Thermo Fisher Scientific | 28025013 |

supplemental table 2

Assignment of lipid molecular species by IMS negative ion mode

| Lipid assignment | $[M-H]^-$ (*m/z*) |
|---|---|
| Palmitic acid | 255.25 |
| Oleic acid | 281.3 |
| Stearic acid | 283.35 |
| Arachidonic acid | 303.3 |
| DHA | 327.33 |
| PE (p18:1/16:0), plasmalogen | 700.6 |
| PE (18:0/16:1) | 716.6 |
| PE (p18:0/20:4), plasmalogen | 750.5 |
| PS (18:0/16:0) | 762.6 |
| PE (18:0/20:4) | 766.5 |
| PE (18:0/22:4) | 794.5 |
| PS (18:0/20:4) | 810.6 |
| PS (18:0/22:6) | 834.6 |
| PI (16:0/20:4) | 857.6 |
| PI (18:1/20:4) | 883.55 |
| PI (18:0/20:4) | 885.77 |
